# Leveraging elastic networks in a coarse-grained brownian simulation framework for protein conformational dynamics

**DOI:** 10.64898/2026.09.22.753665

**Authors:** Domenico Scaramozzino, Enrica Ruggeri, Mariia Feshyna, Byung Ho Lee, Laura Orellana

## Abstract

Protein dynamics is fundamental to understand mechanisms. Although atomistic Molecular Dynamics (MD) remains the gold standard for predicting protein motions, its computational cost limits applications at large scales. Here, we present a coarse-grained (CG) simulation framework that combines Elastic Network Models (ENMs) with implicit-solvent Brownian Dynamics (BD) to efficiently generate protein conformational ensembles from native states. Benchmarking the method on more than 8,500 proteins against large datasets of atomistic MD trajectories and multi-state ensembles from Nucleic Magnetic Resonance (NMR), we show that short ENM-BD simulations can reproduce the residue fluctuation profiles and the dominant protein motions with remarkable accuracy. Correlations of residue fluctuations often exceeded 80%, with many proteins showing agreements above 90%, while Principal Component Analysis (PCA) revealed strong correspondences between the essential motions in the ENM-BD simulations and those observed in MD and NMR ensembles. Following CG-to-all-atom reconstruction and short energy minimization, the ENM-BD conformers also achieve high stereochemical quality, which enables their usage in downstream atomistic applications. Finally, we show that the stochastic dynamics emerging from BD trajectories closely aligns with the normal modes encoded in the underlying elastic network, highlighting that most of the intrinsic dynamics of proteins is already embedded in their native 3D topology.

## Introduction

The classical *sequence-structure-function* paradigm of protein biology has evolved into the more comprehensive *sequence-structure-ensemble-function*, recognizing that proteins exist as ensembles of interconverting states^1^. Advances in experimental techniques, including X-ray crystallography^2^ and cryogenic Electron Microscopy (cryo-EM)^3^, have expanded our ability to explore protein conformational ensembles. Yet, experimental methods remain limited by the systems they can probe and by the particular experimental conditions which may differ from the physiological environment. On the other hand, molecular simulations have now become a “*computational microscope*” to relate protein dynamics and function^4,5^. Atomistic Molecular Dynamics (MD) simulations^6^ remain the gold-standard for sampling protein motions, yet their computational cost has motivated the development of several coarse-grained (CG) approaches^7–10^. These methods have successfully captured relevant processes, ranging from protein folding^11^ to the dynamics of folded^12,13^ and intrinsically disordered^14,15^ proteins, demonstrating that atomistic details are not always necessary.

Among CG approaches, Elastic Network Models (ENMs)^16–18^ represent proteins as networks of beads connected by harmonic springs. Combined with Normal Mode Analysis (NMA)^19^, ENMs efficiently identify collective motions that often closely resemble the conformational changes observed from experiments^20–22^. Numerous ENM variants have been proposed to optimize the spring network and improve the agreement with crystallographic B-factors^23–27^ and conformational changes^28–30^. In particular, the essential-dynamics ENM (edENM) by Orellana et al.^31^ defines two main types of elastic springs to generate normal modes with better agreement with MD: one that accounts for the stronger bonded interactions along the backbone, and the other for describing weaker nonlocal contacts, both fitted to reproduce MD flexibility.

A fundamental limitation of NMA, however, is that the large-amplitude displacements generated by simple linear extrapolation of normal-mode vectors rapidly lead to unrealistic and highly distorted structures^29,30^. To overcome this limitation while retaining the ENM efficiency, ENMs could be used in the context of time-dependent simulations. Using a simple two-parameter ENM potential, Emperador et al.^32^ showed that Brownian Dynamics (BD) simulations can reproduce MD-derived fluctuations, particularly for proteins undergoing large-scale domain and loop motions. More recently, the eBDIMS path-sampling method^33,34^ has been developed, coupling the MD-refined edENM with BD and Dynamic IMportance Sampling (DIMS), to generate transition pathways between two known protein end states. This approach, which has also been recently extended to DNA, RNA, and protein-nucleic acid complexes^35^, has proven valuable for exploring conformational landscapes^36^, elucidating molecular mechanisms^37–39^, and interpreting cryo-EM data^40–42^. However, because the DIMS bias requires two end-state conformations, its applicability remains restricted to systems for which two experimentally resolved conformations are available.

Here, we combine ENMs with unbiased BD (ENM-BD) to sample protein conformational dynamics from single structures. Using a benchmark dataset of ∼2,000 monomeric proteins with atomistic MD trajectories and a complementary dataset of ∼7,000 multi-state ensembles from Nuclear Magnetic Resonance (NMR), we show that ENMs can generate conformers with realistic stereochemistry and can reproduce the residue fluctuations and the essential motions observed in MD and NMR ensembles. ENM-BD can therefore provide a computationally inexpensive method to explore the intrinsic dynamics of proteins, generate physically realistic ensembles that do not rely on linear normal-mode extrapolation, offering a quicker alternative to atomistic simulations.

## Materials and Methods

### Elastic Network Model (ENM)

Protein structures were represented using Elastic Network Models (ENMs), in which C_α_ atoms are modeled as point masses connected by harmonic springs. Two ENM variants were initially considered: the standard Anisotropic Network Model (ANM)^18^ and the MD-refined ENM (edENM)^31^. In the ANM, all pairs of C_α_ atoms with equilibrium distances 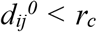 are connected by identical harmonic springs with force constant *k*_*ij*_ = 1 kcal mol^-1^ Å^-2^, using a cutoff distance *r*_*c*_ = 13 Å. In contrast, the edENM introduces a distinction between sequential and spatial interactions, with spring constants defined as:

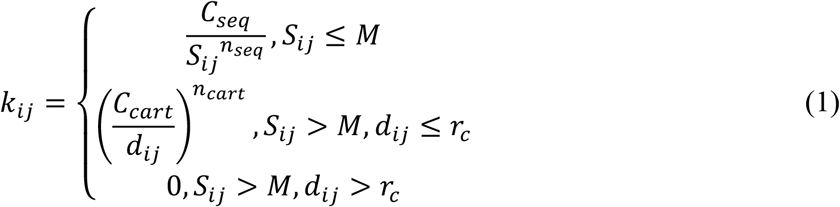

where *S*_*ij*_ is the sequence separation between residues *i* and *j, M* is a sequential cutoff, and *C*_*seq*_ and *C*_*cart*_, and *n*_*seq*_ and *n*_*cart*_ control the strength and distance dependence of sequential and non-local interactions, respectively. The original edENM parametrization was optimized by Orellana et al.^31^ to reproduce MD flexibility using normal modes, corresponding to optimal values: *M* = 3, *r*_*c*_ = 2.9ln(*N*) – 2.9, *N* being the number of residues, *C*_*seq*_ = 60 kcal mol^-1^ Å^-2^, *C*_*cart*_ = 6 kcal^1/6^ mol^-1/6^ Å^2/3^, *n*_*seq*_ = 2, and *n*_*cart*_ = 6. Here, we also introduce a modified variant (edENM^mod^) to improve the protein backbone stability during Brownian Dynamics (BD) simulations. edENM^mod^ keeps the same formulation in Equation 1 but strengthens sequential interactions (*C*_*seq*_ = 600 kcal mol^-1^ Å^-2^) and employs a protein-independent cutoff (*r*_*c*_ = 10 Å).

### Normal Mode Analysis (NMA)

ENMs are normally used in combination with Normal Mode Analysis (NMA)^43^. Within the quadratic approximation, NMA looks for the small-amplitude harmonic oscillations of the system. This is achieved by assembling the 3*N* × 3*N* Hessian matrix ***H***, which contains the second derivatives of the potential energy *V*(***r***) with respect to the chosen coordinate representation. In the case of ENMs, the quadratic potential is simply the elastic energy stored within the spring network^17,18^. Diagonalization of ***H*** yields a complete set of orthonormal eigenvectors and eigenvalues:

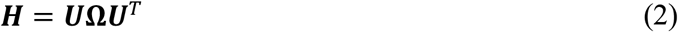

where the columns of ***U*** correspond to the eigenvectors, the normal modes (NMs), and ***Ω*** is the diagonal matrix of eigenvalues, which are related to the mode frequencies^19,44^. From the NMs and their corresponding eigenvalues, root-mean-square fluctuations (RMSFs) related to the thermal oscillations can be derived as^44^:

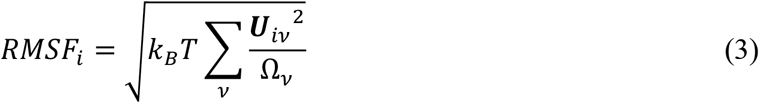

where *k*_*B*_ is the Boltzmann constant, *T* is the absolute temperature, ***U***_***iv***_ is the component of eigenvector *v* at node *i, Ω*_*v*_ the mode eigenvalue, and the summation runs over all non-rigid NMs.

### Brownian Dynamics (BD)

To sample time-dependent conformations and not relying on the linear directionality of normal modes, ENMs were embedded in a Brownian Dynamics (BD) simulation. BD was performed using the Langevin equation of motion^32^:

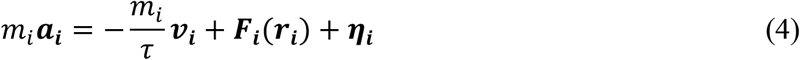

where *m*_*i*_, ***r***_***i***_, ***v***_***i***_, ***a***_***i***_, denote the mass, position, velocity, and acceleration of particle *i*, respectively, *τ* is the friction timescale, ***F***_***i***_ is the deterministic force arising from the ENM potential, and ***η***_***i***_ is a stochastic force accounting for the thermal noise. Assuming a Gaussian white noise and integrating Equation 4 with a Verlet scheme at small time steps *Δt* yields the update equations^32^:

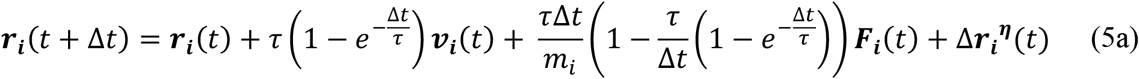

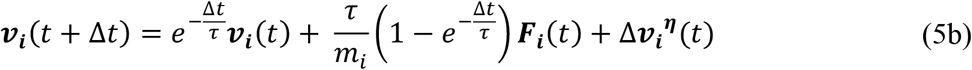

where the stochastic contributions are given by:

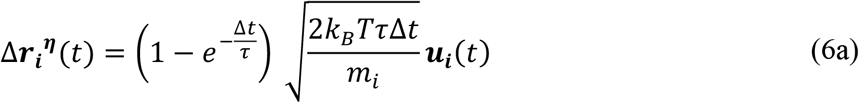

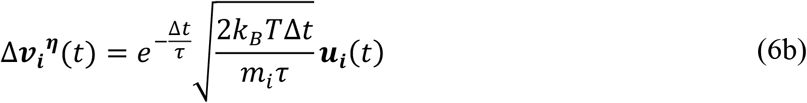

Here, ***u***_***i***_ is a vector of independent Gaussian random variables and *T* the absolute temperature. All BD simulations were performed at *T* = 300 K, using a time step *Δt* = 2 fs and a friction timescale *τ* = 2.5 ps, which is representative of the rotational friction of water^32,45,46^. The initial coordinates 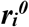 were taken from the Protein Data Bank (PDB) structure, while initial velocities 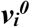 were sampled from a distribution with variance 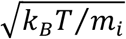. The deterministic force ***F***_***i***_ arises from the ENM potential:

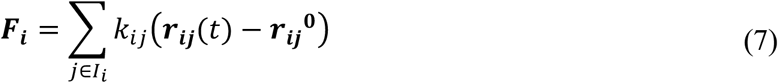

where the sum runs over all interacting neighbors of residue *i*, and 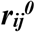 denotes the equilibrium distance in the reference structure. By construction, forces vanish at the initial configuration, which therefore represents a local minimum of the elastic potential^16^. Thermal fluctuations and stochastic forces drive deviations from this state, while the elastic network tends to maintain secondary structures and the overall fold throughout the BD trajectory, but allowing some degree of conformational flexibility.

### Protein Datasets and Benchmarking

To evaluate the sampling performance of the ENM-BD approach, simulations were carried out on two benchmark datasets. The first comprises the 1,938 monomeric proteins from the ATLAS database with MD trajectories^47^. The second consists of 6,704 multi-state ensembles from solution Nuclear Magnetic Resonance (NMR) extracted from the PDB. NMR ensembles were selected to include monomeric proteins with at least 50 residues and a minimum of two deposited NMR models, while excluding entries with missing residues or sequence inconsistencies across models. Ensembles with negligible structural variability (C_α_ root-mean-square deviation, RMSD < 1 Å) were discarded to avoid cases dominated by side-chain rearrangements or ligand-dependent differences.

For MD systems, the initial PDB conformation was used as the starting ENM configuration. For NMR ensembles, the centroid of the ensemble, i.e., the NMR model with minimum RMSD relative to the average coordinates, was selected as the initial seed. For each system, three independent 1 ns-long BD trajectories were generated and combined into a single conformational ensemble. Residue flexibilities were quantified using RMSFs, computed from the aligned positions ***r***_***i***_ as:

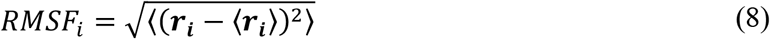

where 〈… 〉 denotes averaging over all conformations in the ensemble.

Principal Component Analysis (PCA)^48–50^ was applied to extract the dominant motions within the ensemble. PCA is a multivariate statistical technique applied to reduce the number of dimensions to describe protein structures and dynamics^48^ and has been widely used to describe the essential motions of proteins from simulation and experimental ensembles^22,33,51^. The input of PCA is an *n* × 3*N* coordinate matrix, ***X***, *n* being the number of structures in the ensemble and *N* the number of residues, usually considering C_α_ atoms. From ***X***, the elements of the covariance matrix, ***C***, are calculated as:

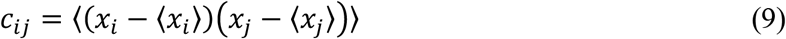

Eigenvalue-eigenvector decomposition is then used to diagonalize ***C***:

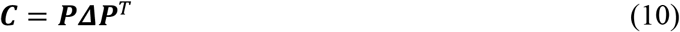

where ***P*** contains the eigenvectors of ***C***, representing the Principal Components (PCs), while the diagonal matrix ***Δ*** contains its eigenvalues, which are related to the amount of variance captured by each PC vector. The agreement between ENM-BD and MD/NMR ensembles was assessed through Pearson Correlation Coefficients (PCCs) of RMSF profiles, as well as the maximum overlaps (O_max_) and root-mean-square inner products (RMSIPs) between the Principal Component (PC) vectors and the dominant subspaces^22^.

We also assessed the integrity of the protein backbone along the simulation by monitoring consecutive C_α_-C_α_ distances, which are expected to remain close to the theoretical value of ∼3.8 Å. To further evaluate the effect of the edENM^mod^ parametrization on the stabilization of local secondary structures in biased BD simulations, the new network parametrization was implemented within eBDIMS2^34^ to generate transition pathways for four benchmark proteins: ribose-binding protein (RBP), RNA endonuclease III (RNaseIII), sarcoplasmic/endoplasmic reticulum Ca^2+^ ATPase1 (SERCA), and the GroEL chaperonin 7-mer^34^. To evaluate the atomistic quality of the generated conformations, C_α_-only ENM-BD conformers were first reconstructed using cg2all^52^ and their atomistic quality was then assessed with MolProbity^53^. In addition, structures reconstructed with atomistic details were subjected to short energy minimizations using the steepest-descent algorithm in GROMACS^54^ (50,000 minimization steps, maximum force at convergence < 1,000 kJ mol^-1^ nm^-1^).

## Results

### ENM-BD can generate stereochemically realistic conformers

A key requirement for our simulations was that the generated conformations remain stereochemically realistic, despite the adopted CG representation. To assess this, we first performed triplicates of 1 ns-long BD simulations for each of the 1,938 monomeric proteins in the ATLAS dataset^47^ starting from the corresponding X-ray structures. We evaluated two existing elastic network parametrizations: the uniform-spring ANM^18^ and the MD-refined edENM^31^. Because these force fields were originally developed for NMA calculations and not in the context of time-dependent simulations, their capability to maintain realistic protein geometries along non-linear trajectories had not been previously checked.

BD simulations with the uniform-spring ANM were found to produce substantial distortions of the CG backbone, with distances between consecutive C_α_ atoms spanning from below 3.0 Å to above 4.5 Å (Fig. 1A). Replacing the ANM with the MD-refined edENM (which employes stronger interactions for sequential backbone neighbors) reduced these distortions and produced a much narrower distance distribution centered at the expected C_α_-C_α_ values (∼3.8 Å). However, a residual difference between the edENM and the MD C_α_-C_α_ distributions remained (Fig. 1A). To address this, we introduced a modified edENM (edENM^mod^; see Methods), which increases the strength of sequential interactions by a factor of 10. This simple modification was able to yield C_α_-C_α_ values virtually indistinguishable from those observed in the conformations generated from atomistic MD (Fig. 1A and Supplementary Table 1).

**Figure 1.**
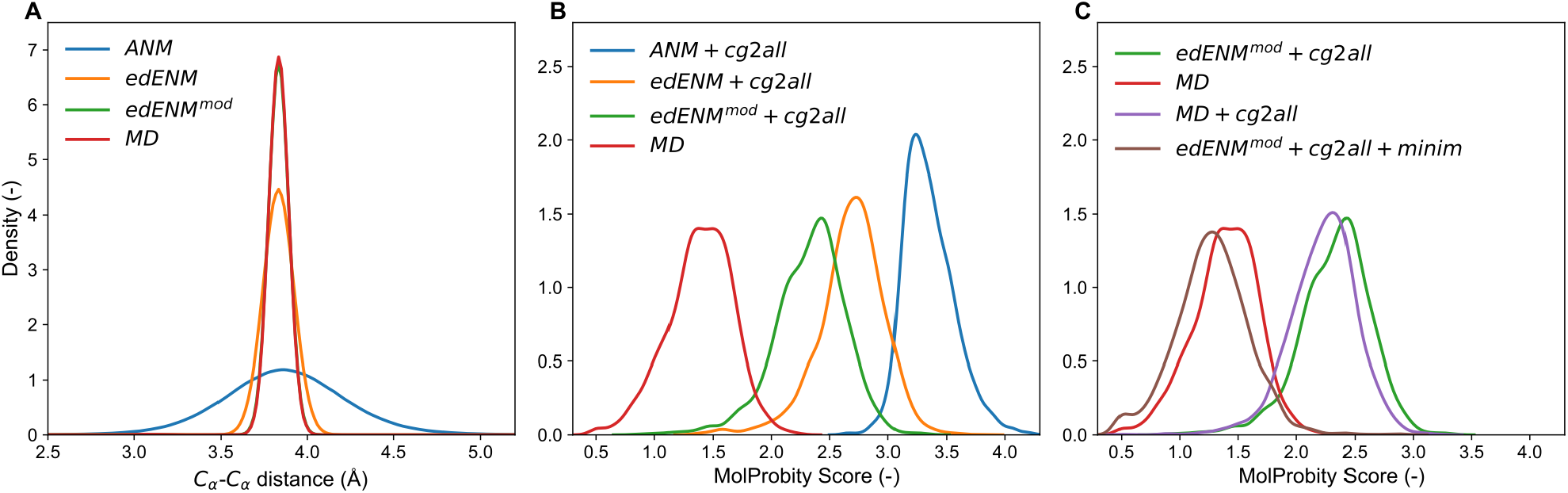
Stereochemical quality of ENM-BD conformers evaluated across the ATLAS dataset. (A) Distribution of consecutive C_α_-C_α_ distances in atomistic MD (red) and ENM-BD simulations carried out using ANM (blue), edENM (orange), and edENM^mod^ (green). The stronger sequential interactions of edENM^mod^ preserve the backbone geometry. (B) MolProbity scores of atomistic structures reconstructed with cg2all from ENM-BD conformers compared with the corresponding MD frames. Progressive refinement of the elastic network improves the stereochemical quality of the generated conformers. (C) MolProbity scores of atomistic MD conformers (red) and edENM^mod^-BD conformers after cg2all reconstruction (green) as in panel (B), compared with MD conformers stripped to C_α_-only atoms and subsequently reconstructed with cg2all (purple), as well as atomistic edENM^mod^ conformers following a short energy minimization (brown). The deterioration in stereochemical quality primarily arises from the atomistic reconstruction and can be easily recovered with short energy minimizations.

To verify that the improved local geometry could be maintained also in larger-amplitude conformational transitions, we incorporated edENM^mod^ into eBDIMS2 and simulated large-scale transition pathways between open and closed conformations of four proteins that we have previously investigated in detail^34^. As expected, edENM^mod^ was found to improve substantially the quality of the CG bakbone compared to the previous eBDIMS implementations (Supplementary Fig. 1), while preserving the overall motion trajectories (Supplementary Fig. 2). These results confirm that strengthening the ENM backbone interactions improves the stereochemical quality not only in unbiased simulations, but also in larger-scale transitions between end-state conformers.

The atomistic quality of the conformers was found to mirror the backbone quality at the CG level. Conformations generated with ANM generally exhibited poorer stereochemistry, as reflected by relatively high MolProbity scores (Fig. 1B). Progressively refining the elastic network improved the quality of the reconstructed conformations, although their MolProbity scores remained worse than those from atomistic MD (Fig. 1B and Supplementary Table 2). To determine whether this residual difference might originate from the atomistic reconstruction itself^52^, we extracted frames only with C_α_ atoms from the MD simulation and subsequently reconstruct them with cg2all (“MD + cg2all” in Fig. 1C). This was in fact found to exhibit a marked deterioration in the stereochemical quality of the MD-generated conformations, leading to MolProbity scores comparable to those obtained for edENM^mod^ conformers (Fig. 1C), suggesting that the quality gap between edENM^mod^-BD and MD is mainly attributable to inherent limitations of reconstruction procedures. Finally, we evaluated whether the reconstructed edENM^mod^-BD conformations could recover better atomistic quality through short energy minimizations. As expected, this simple refinement increased the stereochemical quality of the BD conformers to a level indistinguishable from that of atomistic MD (Fig. 1C). Importantly, these minimizations preserve the overall CG conformation (RMSD_minim_ < 0.5 Å), mostly re-optimizing side-chain orientations and relieving steric clashes induced by the atomistic reconstruction (Supplementary Fig. 3). Taken together, these results demonstrate that ENM-BD can be used to generate new protein conformations with realistic stereochemistry, not only at the CG but also at the atomistic level.

### How does ENM-BD sampling compare with atomistic MD?

To evaluate how ENM-BD simulations compare with the dynamics obtained by atomistic MD simulations, we looked again at the ensembles generated with short (1 ns) BD simulations for all 1,938 proteins in the ATLAS dataset. Residue fluctuation profiles (RMSFs) obtained from the ENM-BD simulations were found to exhibit generally good agreements with the longer, 100 ns MD ensembles, with average Pearson Correlation Coefficients (PCCs) exceeding 75% across the entire dataset (Supplementary Table 3). The refined elastic networks slightly outperformed the uniform-spring ANM (Fig. 2A). In particular, edENM^mod^ achieved RMSF correlations above 80% for 1,133 proteins (∼58% of the dataset), and for 599 proteins (∼30%) it exhibited correlations even exceeding 90%, reaching values as high as ∼99%.

**Figure 2.**
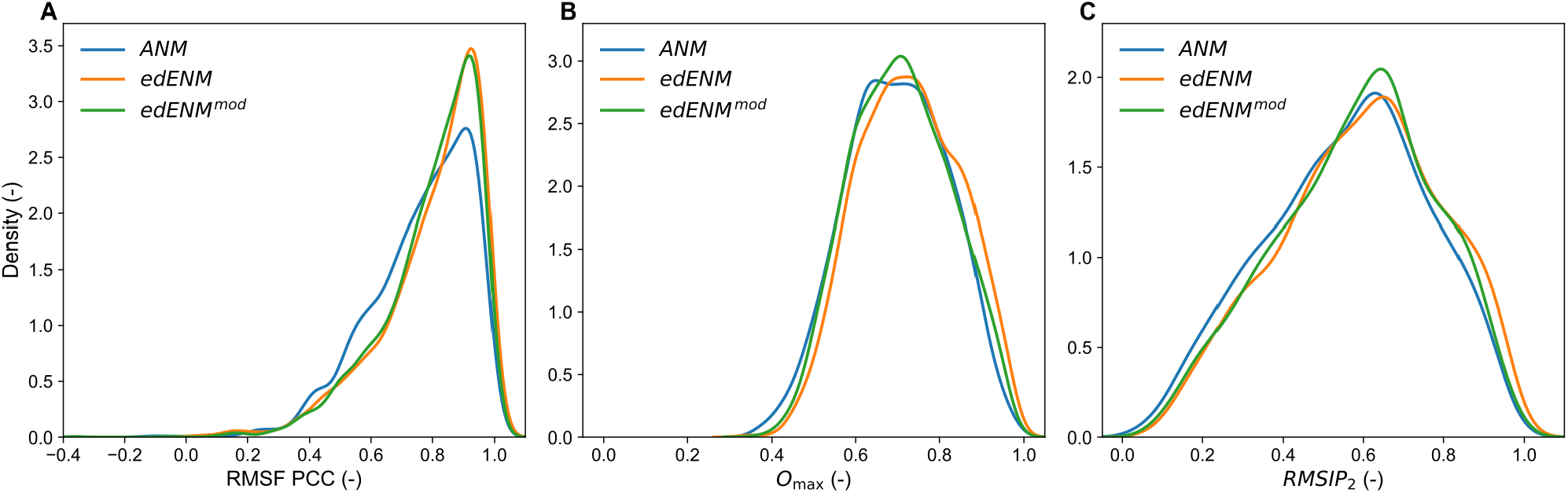
ENM-BD reproduces relative fluctuations and essential motions from atomistic MD. (A) Distribution of Pearson Correlation Coefficients (PCCs) between C_α_ RMSFs obtained from MD and ENM-BD simulations using ANM (blue), edENM (orange), and edENM^mod^ (green). (B) Distribution of maximum overlap (Omax) between the ten leading PCs of MD and ENM-BD conformational ensembles. (C) Distribution of RMSIP between the essential subspaces defined by the two leading PCs (RMSIP2) from MD and ENM-BD ensembles.

While the RMSF comparison allows to quantify the agreements in terms of relative amplitude of residue fluctuations, it does not provide information on the motion directionalities. Thus, we next compared the essential motion directions observed within ENM-BD and MD ensembles using Principal Component Analysis (PCA). The similarity of the sets of dominant motions was assessed by computing the maximum overlap (O_max_) between the ten lowest Principal Components (PCs) and comparing the two essential subspaces using root mean squared inner products (RMSIPs; see Methods). Across the whole ATLAS dataset, O_max_ values were on average ∼70% (Supplementary Table 3), with numerous proteins exhibiting maximum vector overlaps above 80-90% (Fig. 2B). The similarity between the two-dimensional essential subspaces (RMSIP_2_) was on average ∼60%, with a substantial fraction of proteins displaying RMSIPs greater than 80% (Fig. 2C). Taken together, these results demonstrate that short ENM-BD simulations are not only able to capture with satisfactory accuracy the relative amplitudes of residue fluctuations, but also the 3D motion directionalities in the essential motions encoded in longer, atomistic MD.

Focusing on the BD simulations performed with the edENM^mod^ variant, we found that 184 proteins achieved performance values higher than 80% for all three metrics (RMSF PCC, O_max_, and RMSIP_2_), and 19 proteins even exceeded 90% in all metrics. Two of such proteins are shown in Fig. 3. The upper panels illustrate the HR-1 domain of the *D. Melanogaster* caprin homolog. ENM-BD reproduced the MD residue fluctuations with a PCC of 92% (Fig. 3A), and the first three principal components exhibited an almost perfect one-to-one correspondence (Fig. 3B), with O_max_ = 93% and RMSIP_2_ = 91%. Notably, the variances captured by the dominant PCs were also consistent between the two ensembles (Supplementary Fig. 4A). Despite the ENM-BD simulations employ shorter sampling timescales, the two ensembles also showed similar values of structural deviations (RMSD) from the initial X-ray model (Fig. 3C). Another interesting case is the *L. pneumophila* DUF5617 domain-containing protein (Fig. 3; lower panels). Here, RMSF profiles exhibited a PCC of 98% (Fig. 3D), and the dominant PC vectors were also nearly identical (O_max_ = 97%, RMSIP_2_ = 95%; Fig. 3E), again with matching variance profiles (Supplementary Fig. 4B). In this case, ENM-BD produced lower deviations from the initial structure, with a maximum RMSD of ∼11 Å, compared with the ∼16 Å for the MD ensemble (Fig. 3F). This somewhat lower sampling is also reflected by lower ENM-BD RMSFs compared to MD (Fig. 3D). These results suggest that short ENM-BD simulations can thus be used to reproduce the essential directions of motions as well as the relative fluctuation profiles observed in longer MD simulations. Absolute fluctuation amplitudes can be quantitatively reproduced only in some cases, and not in others (additional details below).

**Figure 3.**
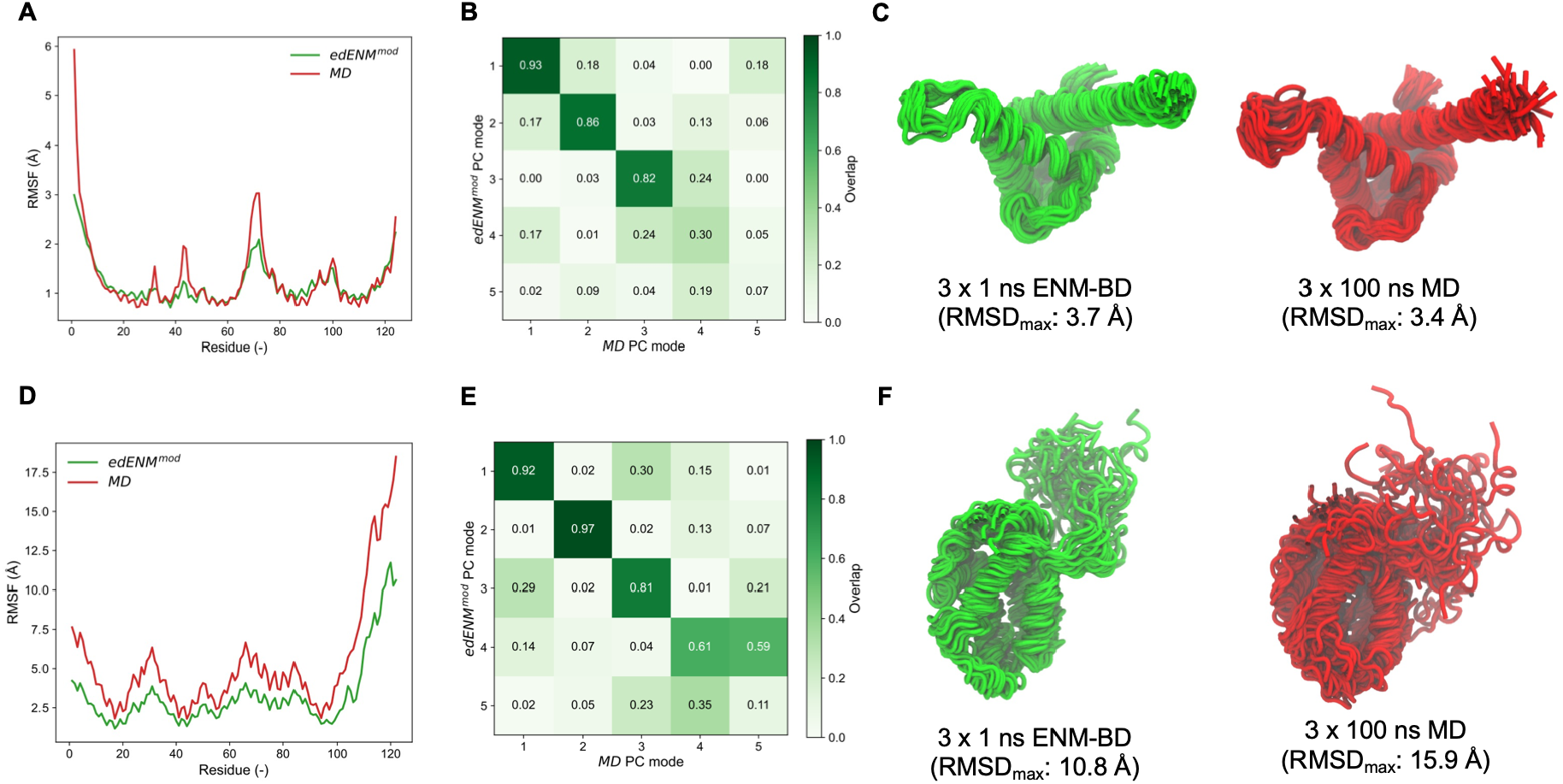
Examples of ENM-BD ensembles with high agreement with atomistic MD. (A-C) HR-1 domain of the *D. Melanogaster* caprin homolog (PDB: 6bk4, chain: A); (D-F) DUF5617 domain-containing protein from *L. pneumophila* (PDB: 4wrp, chain: A). Panels (A) and (D) compare C_α_ RMSFs obtained from ENM-BD (green) and MD (red). Panels (B) and (E) show overlap matrices between the five leading PCs from ENM-BD and MD ensembles, with overlap values ranging from 0 (white) to 1 (dark green). Panels (C) and (F) display C_α_-tube representations of the conformational ensembles sampled by 1 ns-long ENM-BD (green) and 100 ns-long MD (red). Structures were rendered with VMD^55^.

We also looked for cases of poor correlations. The only protein in the entire ATLAS dataset exhibiting extremely poor scores (< 40%) in all three metrics (RMSF PCC, O_max_, and RMSIP_2_) was the 571-residue methanol dehydrogenase subunit 1 from *M. Methylotrophus* (Supplementary Fig. 5). Here, the ENM-BD simulations remained extremely close to the initial X-ray structure (RMSD < 1 Å) and failed to reproduce the large RMSF peaks associated to localized loop motions observed in MD (RMSF PCC ∼15%). Notably, running longer, 100 ns ENM-BD simulations (consistent with the ATLAS MD timescale) did not enhance the sampling (Supplementary Fig. 6A-B). The essential dynamics analysis showed that ENM-BD trajectories were mostly spread over multiple, low-variance PC modes, with no meaningful overlaps with the essential motions from MD (Supplementary Fig. 5). While this was the only case in the ATLAS dataset where the ENM-BD achieved extremely poor agreement with MD, this is informative for highlighting an important difference between ENM-BD and atomistic MD. In the ENM, particle-particle interactions are only defined by harmonic potentials centered on the initial structure. In this way, the spring contacts between two residues tend to stabilize their distance observed in the starting topology^16^, as long as they do not vanish due to the random stochastic motion. In contrast, MD force fields include many environment-dependent interactions that often lead to quick departures from the initial crystal structure in solution. For example, in the case of methanol dehydrogenase, the loop at residues 420-425 can undergo a large conformational rearrangement in MD (RMSF ∼ 7 Å), whereas in ENM-BD the same region remains strongly confined to thermal fluctuations (∼ 1 Å), due to the constraining effect of the initial, extremely compact elastic network topology (Supplementary Fig. 5).

We observed that running longer ENM-BD simulations does not generally lead to an increase in the sampling of larger-scale dynamics (Supplementary Fig. 6). As mentioned above, for proteins with compact topologies the constraining effect of the initial network makes it hard for springs to break or new springs to form, so that running longer ENM-BD simulations lead to almost no gains in sampling new conformations (Supplementary Fig. 6A-B). Proteins with more loosely packed topologies or floppy loops and tails might benefit for longer BD timescales to generate additional conformations (Supplementary Fig. 6D). However, this will mostly affect the flexible parts, not leading to a dramatic change in the assessment of the overall conformational flexibility (Supplementary Fig. 6C). These observations suggest that most of the ENM-BD conformational space can be recovered early during the trajectories and that running longer BD simulations will not dramatically change the sampling outcomes.

To compare the amplitude of sampling of short ENM-BD simulations against the longer MD trajectories in the ATLAS dataset, we computed maximum C_α_ RMSD values for all ATLAS proteins from their initial X-ray structure. As shown in Fig. 4A, the 1 ns-long ENM-BD simulations had in fact a tendency to sample smaller conformational changes compared to the 100 ns MD, confirming a more restricted exploration of the conformational space. On average, the 100 ns-long MD trajectories reached maximum RMSDs of ∼6.4 Å, whereas 1 ns-long ENM-BD exhibited lower sampling values of ∼2.0 Å (ANM), ∼4.7 Å (edENM), and ∼3.3 Å (edENM^mod^). As expected, ANM exhibited the most rigid behavior. This is mostly due to the uniform spring constants (1 kcal mol^-1^ Å^-2^) adopted in the ANM: while these are too weak for describing local contacts, which leads to poor stereochemistry, they get overly strong for describing nonlocal interactions, which results in a system that becomes too rigid. Due to the differentiation between stronger local contacts and weaker nonlocal interactions, edENM enabled a larger-scale conformational sampling (Fig. 4A). The edENM^mod^ variant, which strengthens local contacts with the goal of improving the stereochemical quality of the conformations (Fig. 1), now pays the prize becoming slightly more rigid than edENM. Yet, it can still sample more heterogeneous conformations compared to ANM (Fig. 4A). In the remaining of the text, we will only focus on the results obtained with edENM^mod^.

**Figure 4.**
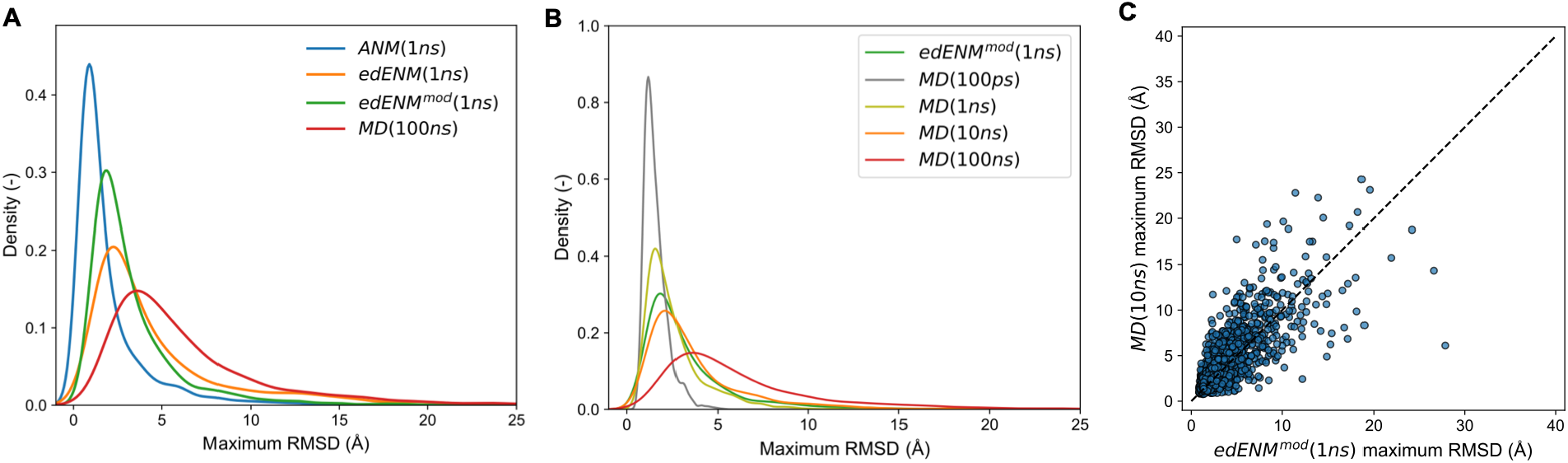
Comparison between ENM-BD and MD sampling amplitude for the ATLAS dataset. (A) Maximum C_α_ RMSD from the initial X-ray structure sampled with 100 ns MD (red) and shorter, 1ns-long ENM-BD simulations (ANM: blue, edENM: orange, and edENM^mod^: green); (B) Comparison of maximum RMSD values between edENM^mod^-BD simulations (1 ns) and MD at different timescales (100 ps, 1 ns, 10 ns, and 100 ns); (C) Scatter-plot comparison between 1 ns-long edENM^mod^-BD and MD sampling after 10 ns, with each point representing a protein in the ATLAS dataset. 1 ns edENM^mod^-BD simulations sample more diverse conformations than 1 ns atomistic MD.

To evaluate how short BD sampling compares with MD simulations at different timescales, we then computed maximum RMSDs obtained in the ATLAS trajectories after the first 100 ps, 1 ns, 10 ns, and 100 ns (Fig. 4B). RMSD values at these MD timescales averaged ∼1.6 Å for 100 ps, ∼2.6 Å for 1 ns, ∼3.8 Å for 10 ns, and ∼6.4 Å for 100 ns. Comparing with the sampling from 1 ns-long edENM^mod^-BD (average RMSD ∼3.3 Å), these results suggest that 1 ns BD samples on average more than 1 ns MD (Fig. 4B and Supplementary Fig. 5), and that the amount of sampling achieved by 1 ns BD simulations is actually more comparable to performing 10 ns of atomistic MD (Fig. 4C).

We must stress here that comparing these timescales does not necessarily have a physical meaning. First, because by adopting a CG model (with implicit solvent) we remove many of the fast degrees of freedom that are crucial in MD simulations. Secondly because, as already highlighted above, longer ENM-BD simulations do not necessarily increase conformational sampling, unlike MD which can capture rare events if performed at longer, microsecond timescales. This comparison is meant to put the short BD simulations in the (more familiar) context of MD timescales, suggesting that when we run 1 ns ENM-BD we can expect to obtain an amount of RMSD sampling which is typically retrieved in ∼10 ns atomistic MD. Yet, at a fraction of the computing cost (see below).

### ENM-BD sampling agrees with multi-state ensembles from NMR

Motivated by earlier observations that MD essential subspaces agree with multi-state ensembles that fit NMR data^56^ and by the fact such NMR ensembles have been benchmarked against ENM normal modes^21,57,58^ and other CG simulation methods^59,60^, we next performed ENM-BD simulations on a benchmark of 6,704 protein ensembles in the PDB obtained from solution NMR experiments. For these simulations, we only employed the refined edENM^mod^ parametrization and performed, again, triplicates of 1 ns-long BD.

The stereochemical quality of the generated conformers mirrored what already observed for the MD benchmark. Consecutive C_α_-C_α_ distances remained centered around the expected value of ∼ 3.8 Å (Supplementary Table 4, Supplementary Fig. 6A), confirming that edENM^mod^ preserves the CG backbone geometry also when starting from NMR models. As for the MD dataset, the atomistic reconstructions from the ENM-BD ensembles initially exhibited suboptimal MolProbity scores. Also in these cases, atomistic stereochemistry could be restored to higher quality by means of short energy minimizations (Supplementary Table 5, Supplementary Fig. 6B). Interestingly, deposited NMR models exhibited, on average, poorer atomistic stereochemistry than the X-ray structures from the ATLAS dataset. This is consistent with previous observations that NMR models generally display low quality compared to structures from X-ray crystallography^61^. In this case, reconstructing NMR conformations from their C_α_ traces with cg2all tends to produce better models compared to the deposited structures (Supplementary Table 5, Supplementary Fig. 6B).

We next evaluated the ability of ENM-BD to reproduce the conformational variability observed in these multi-state ensembles from solution NMR. Overall, agreement metrics were even higher than those obtained for the ATLAS dataset (Fig. 5), indicating that the ENM-BD ensembles are also consistent with the dynamics indirectly captured from NMR data. RMSF profiles exhibited an average PCC exceeding 80% (Fig. 5A, Supplementary Table 6). Across the dataset, 4,459 proteins (∼67% of the dataset) showed RMSF correlations above 80%, while 2,916 (∼43%) exceeded even 90%, with best-performing cases approaching nearly perfect RMSF agreement (∼99%). As observed for the ATLAS dataset, the agreement was also found when looking at the directionality of the essential (PCA) motions in the ensemble. Maximum overlaps between PC vectors averaged ∼70% (Supplementary Table 6, Fig. 5B), with 1,607 proteins exhibiting values above 80% and 310 proteins above 90%. The similarity between the dominant PC subspaces (RMSIP_2_) was on average ∼60% (Supplementary Table 6, Fig. 5C), exceeding 80% for 1,162 proteins and 90% for 220 proteins. Together, these results demonstrate that ENM-BD can reproduce the relative amplitude of residue fluctuations and the essential motions extracted from multi-state NMR ensembles. Interestingly, unlike the comparison with atomistic MD (Fig. 4A), short ENM-BD simulations enabled to sample amplitudes of motions with a good quantitative match with those observed in NMR, resulting in comparable distributions of maximum RMSD values between the ENM-BD and NMR ensembles (Fig. 5D).

**Figure 5.**
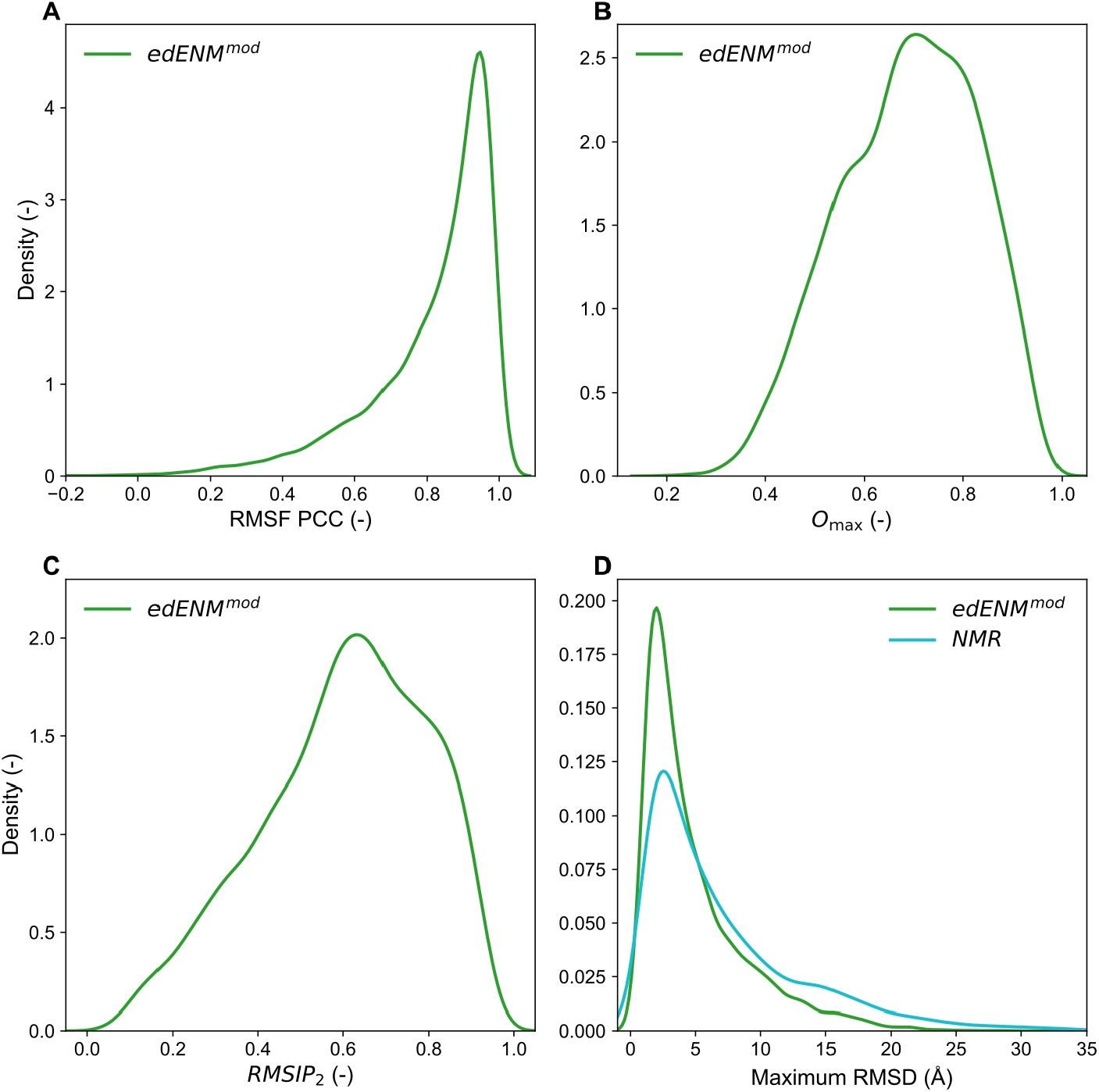
ENM-BD performance evaluated across the NMR dataset. (A) Correlation between C_α_ RMSFs computed from the NMR ensembles and ENM-BD simulations. (B) Maximum overlaps (Omax) between PC vectors derived from NMR and ENM-BD ensembles. (C) RMSIP_2_ between NMR and BD ensembles. (D) Comparison of maximum C_α_ RMSD values within NMR ensembles (cyan) and those sampled by ENM-BD simulations (green).

Across the 6,704 NMR ensembles, 740 proteins were found to exhibit values > 80% for all performance metrics (RMSF PCC, O_max_, and RMSIP_2_). Remarkably, 92 of these proteins even exceeded 90% in all metrics, indicating an almost complete correspondence between ENM-BD and NMR dynamics. Figure 6 shows four of these examples, with proteins spanning an array of different folds. In all four cases, the variance captured by the principal components was also consistent between the two ensembles (Supplementary Fig. 7). The C-terminal domain of human Lamin-B2 exhibited good agreement between ENM-BD and NMR, with an RMSF PCC of 95% and nearly identical essential motions (O_1,1_ = 93%, O_2,2_ = 96%, RMSIP_2_ = 96%; Fig. 6A). Likewise, the synthetic ferredoxin-like protein shown in Fig. 6B displayed highly correlated fluctuation profiles (PCC = 96%) and almost indistinguishable dominant motions (O_1,1_ = 94%, O_2,2_ = 89%, RMSIP_2_ = 93%). Similar levels of agreement were observed for the fibronectin type III domain of human tenascin X (Fig. 6C) and DUF971 domain-containing protein from *Pseudomonas Aeruginosa* (Fig. 6D), with RMSF correlations of 94% and 95%, respectively, and maximum PC overlaps reaching 96% and 95%. In these two latter cases, the first two PC modes were swapped between the ENM-BD and NMR ensembles, while the corresponding subspaces remained highly correlated (RMSIP_2_ = 96% and 93%, respectively).

**Figure 6.**
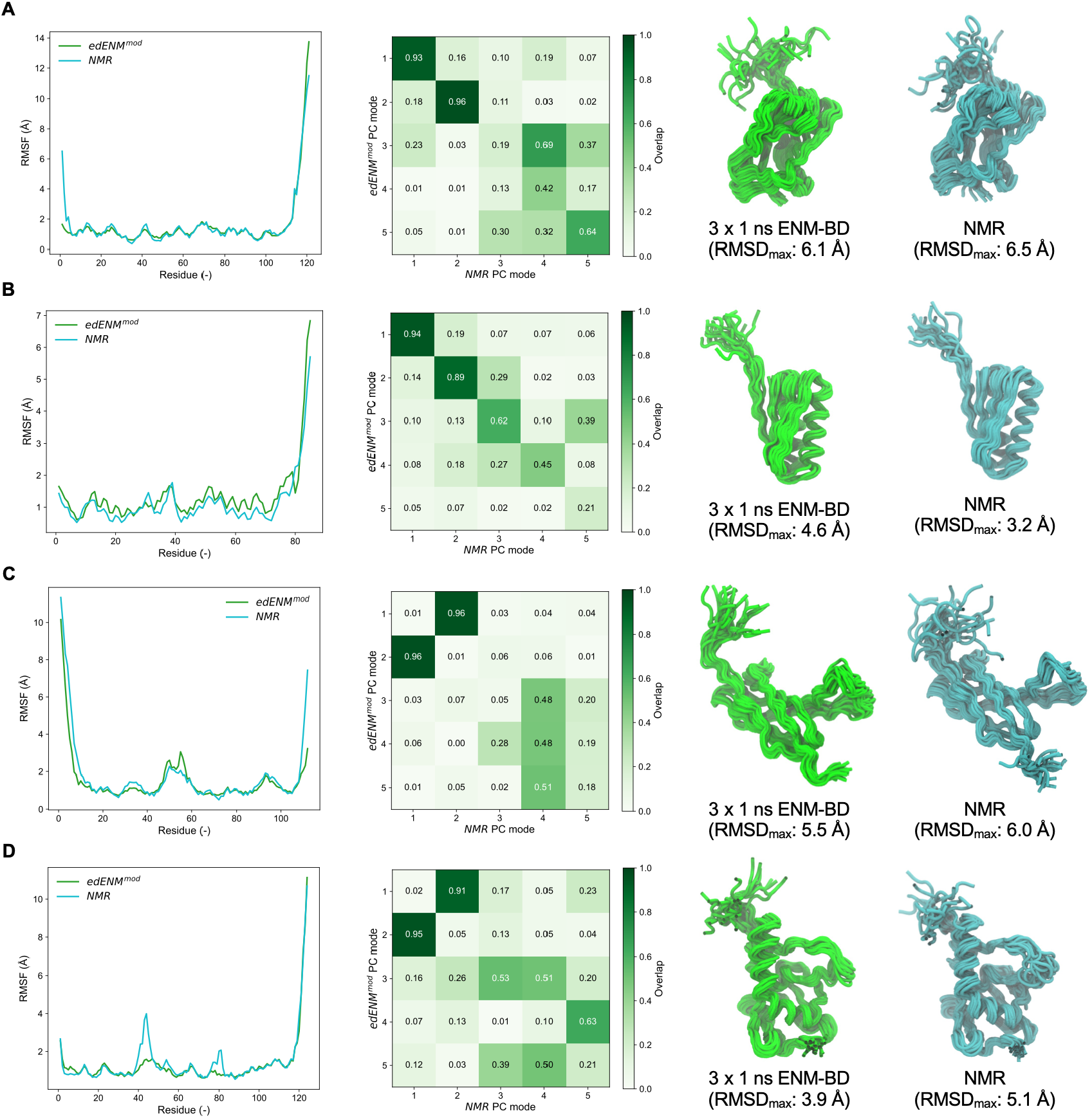
ENM-BD conformational ensembles with high agreement with NMR. (A) C-terminal domain of *H. Sapiens* Lamin-B2 (PDB: 2lll, chain: A); (B) synthetic *de novo* design of ferredoxin-like fold protein (2kl8, A); (C) fibronectin type III domain of *H. Sapiens* tenascin X (2cui, A); (D) DUF971 domain-containing protein from *P. Aeruginosa* (2l6p, A). Left panels compare C_α_ RMSF profiles obtained from ENM-BD (green) and NMR ensembles (cyan). Middle panels show overlap matrices between the five leading principal components of the ENM-BD and NMR ensembles. Right panels display C_α_-tube representations of the conformational ensembles by ENM-BD (green) and NMR (cyan).

To identify limitations of the ENM-BD framework to describe the dynamics observed from the NMR ensembles, we also examined the worst-performing cases in this benchmark. Remarkably, only two of the 6,704 proteins exhibited values below 30% for all performance metrics (Supplementary Fig. 8): the proteinase inhibitor IIA from *Bos Taurus* (PDB: 1bus; PCC = 23%, RMSIP_2_ = 16%) and the human vesicle-associated membrane protein-associated protein B/C (PDB: 2mdk; PCC = −10%, RMSIP_2_ = 15%). Unlike the lowest-performing example identified in the ATLAS benchmark (Supplementary Fig. 5), neither protein displays obvious structural features, such as an unusually compact structure or a highly constrained topology, that would hinder ENM-BD sampling. However, in both cases, NMR ensembles were observed to comprise a relatively small number of deposited conformers (only 5 for 1bus and 10 for 2mdk), fewer than the ∼20 models typically deposited in NMR entries. Moreover, the NMR fluctuation profiles, particularly for 2mdk, are less smooth than those observed in other cases (Fig. 6), suggesting that poor agreement may also result from limitations in the dynamics observed from multi-state ensembles from NMR data.

### ENM-BD spontaneously follows the intrinsic motions embedded in the normal modes

ENMs have mostly been used in combination with NMA^17,18,43^, where collective motions are described as linear combinations of low-frequency normal modes (NMs). Because ENM-BD samples protein motions through stochastic dynamics without explicitly following NM vector directions, we investigated whether the resulting BD trajectories reflect the intrinsic NM motions embedded in the underlying ENM. To this end, we performed NMA using edENM^mod^ for every protein in the ATLAS and NMR datasets and compared the resulting RMSF profiles and the dominant low-frequency NMs against the essential PC modes obtained from ENM-BD simulations.

We found that ENM-BD and ENM-NMA exhibited an almost complete agreement. RMSF profiles derived from the two approaches showed an average PCC exceeding 93% (Supplementary Table 7). Likewise, the dominant motions sampled by BD aligned with the lowest-frequency NMs, with average O_max_ values above 90% and RMSIP_2_ exceeding 80% (Supplementary Table 7). This close correspondence demonstrates that BD naturally follows the intrinsic NMA dynamics encoded within the elastic network, despite being based on a stochastic simulation with implicit solvent and not relying on the linear approximation underlying NMA. As a result of the high agreement between ENM-BD and ENM-NMA dominant dynamics, ENM-BD and ENM-NMA approaches displayed nearly identical performance metrics when benchmarked against the MD trajectories (Fig. 7A-C) and NMR ensembles (Fig. 7D-F).

**Figure 7.**
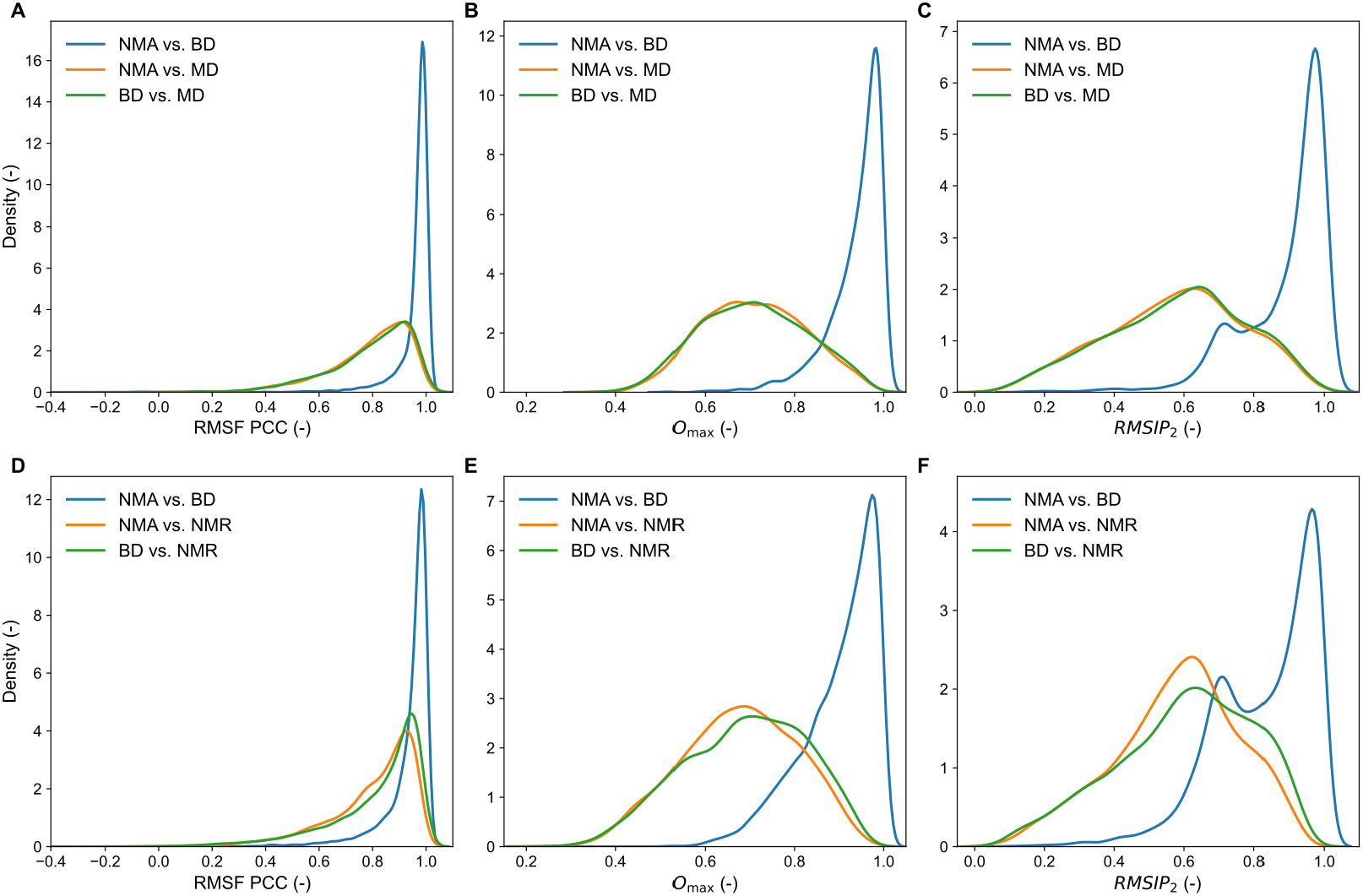
ENM-BD trajectories naturally follow normal modes. (A-C) Comparison between classical ENM-NMA and ENM-BD simulations against the ATLAS dataset. (D-F) Same benchmark against the NMR dataset. Panels (A) and (D) show PCC values between RMSF profiles. Panels (B) and (E) report O_max_ values between NM vectors and PC vectors from ENM-BD, MD, NMR ensembles. Panels (C) and (F) display RMSIP_2_ values. Blue distributions compare ENM-BD with ENM-NMA, orange distributions compare ENM-NMA with MD or NMR, and green distributions compare ENM-BD with MD or NMR.

These results show that ENM-BD tends to closely follow the intrinsic motions encoded in the ENM, despite not relying on explicit NMA calculations, suggesting that the dominant motions described by the lowest-frequency NMs also naturally emerge from stochastic BD trajectories. At the same time, ENM-BD allows to overcome the limitations of NMA. As shown above, BD simulations generate stereochemically realistic conformations up to several Å from the starting structure (Supplementary Fig. 3), avoiding the severe structural distortions that arise by linear NM extrapolation^29,30^. Rather than producing displacements along a fixed set of linear vectors, ENM-BD explores the conformational landscape through stochastic dynamics, allowing broader conformational sampling (Fig. 8). Moreover, because BD is intrinsically stochastic, independent replicas naturally produce distinct BD trajectories, leading to a broader exploration of conformational spaces in different directions (Supplementary Fig. 9). Together, these results suggest that ENM-BD can be seen as a natural extension of ENM-NMA, preserving the ability to identify intrinsic protein motions embedded in the 3D native fold as with NMA, while providing a simulation-based approach to generate broader and more realistic conformational ensembles (Fig. 8).

**Figure 8.**
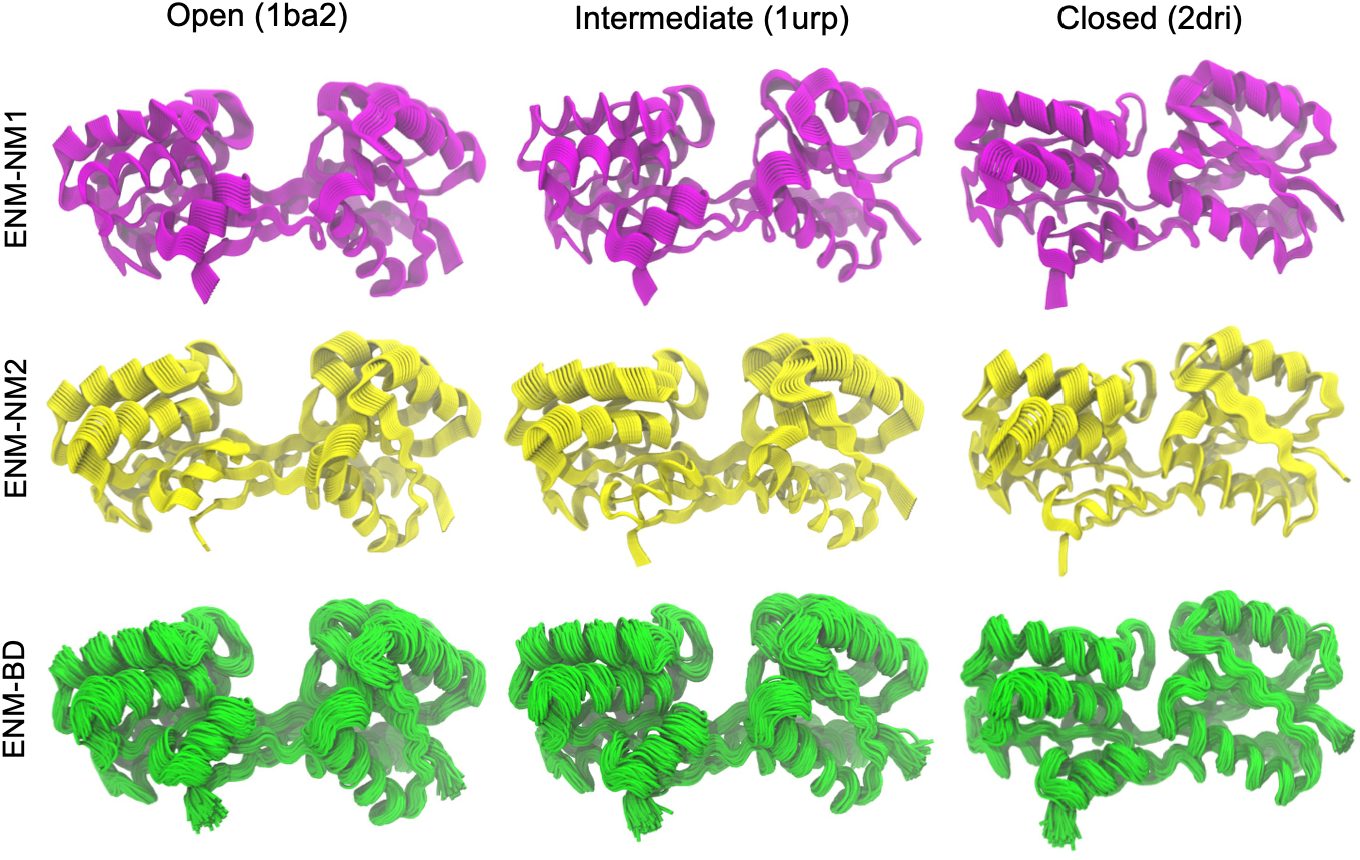
ENM-BD generates richer conformational ensembles than linear normal modes. The first and second rows show conformational ensembles generated for ribose-binding protein (RBP) by linear extrapolations along the first (NM1, magenta) and second (NM2, yellow) low-frequency normal modes, respectively. The third row displays the ensemble of conformations obtained from three independent 1 ns BD simulations. Columns correspond to calculations initiated from the open (PDB: 1ba2), intermediate (1urp), and closed (2dri) conformation from X-ray experiments. Projections of the conformations onto the PCA space derived from experimental RBP structures^33,34^ are shown in Supplementary Fig. 9.

## Discussion

Proteins do not exist as static structures but rather populate ensembles of distinct conformations. Following the success of AlphaFold in predicting single protein structures^62,63^, the next challenge is to generate conformational ensembles^64–66^, which is important to elucidate allosteric mechanisms^67^ and boost allosteric drug discovery^68^. Although several machine-learning approaches have recently been proposed to predict structural ensembles directly from sequences^69–74^, atomistic MD remains the reference approach because protein motions emerging from physically-grounded force fields can provide mechanistic explanations. Yet, despite substantial advances in computational power and the growing availability of large MD databases^47,75,76^, generating ensembles for large protein datasets remains computationally prohibitive. CG approaches therefore represent an attractive alternative for exploring protein dynamics at large scales, as recently demonstrated by the possibility to generate ensembles of intrinsically disordered regions of the entire human proteome^15^.

Here, we demonstrate that coupling ENMs with an implicit-solvent BD framework can provide a simple yet effective approach to extract information on protein flexibility, its dominant motions, and generate physically realistic conformations for large protein datasets. Traditionally, ENMs have been used in combination of NMA to predict directions of conformational changes^20^. This however poses problems when generating new conformers following linear mode extrapolations since severe structural distortions accumulate quickly^29,30^. ENM-BD instead produces time-dependent, non-linear, stochastic trajectories which do not rely on linear vector directions. By introducing a simple modification on the previously developed edENM force field^31^, we show that the resulting CG conformations can be reconstructed with full atomistic detail, achieving atomistic quality comparable to that of MD-derived structures (Fig. 1).

We benchmarked ENM-BD on more than 8,500 monomeric proteins against large-scale datasets of atomistic MD trajectories^47^ and solution NMR ensembles from the PDB. Despite the adopted minimalistic representation, ENM-BD simulations were able to reproduce the conformational dynamics of both reference dataset with high agreement (Figs. 2 and 5). Residue fluctuation profiles exhibited average correlations of ∼80%, with a substantial fraction of proteins reaching correlations above 90%. Likewise, the dominant motions extracted by PCA showed strong agreement, with average similarities of ∼70% for individual motion vectors and ∼60% for the corresponding essential subspaces, and numerous proteins displaying an almost one-to-one correspondence between the principal motions sampled by ENM-BD and those observed in MD (Fig. 3) or NMR (Fig. 6) ensembles.

A major advantage of the ENM-BD framework is its computational efficiency. The ATLAS dataset comprises 1,938 monomeric proteins spanning sizes from 38 residues (*E. Coli* translation initiation factor 4E-binding protein 1) to 2,128 residues (*P. Luminescens* TcdB2). Performing three independent and sequential 1 ns ENM-BD replicas for the entire dataset required approximately 2.5 days of wall-clock time on a standard Linux workstation (Intel^*®*^ Core i9-13900K, 16 CPU threads with OpenMP parallelization). At the level of individual trajectories, this corresponds to an average runtime of ∼37 seconds per replica, ranging from ∼20 s for the smallest proteins to ∼3.5 min for the largest TcdB2 (Fig. 9). Note that this amount of BD sampling (1 ns) is enough to obtain good correlations with MD flexibility and motion directionality (Fig. 2), while simulating for longer times do not necessarily lead to increased sampling (Supplementary Fig. 6). We also showed that 1 ns BD sampling roughly corresponds, if measured in terms of maximum RMSD from the starting structure, to ∼10 ns atomistic MD (Fig. 4), which however require many more CPU-hours than those utilized here. As shown in Fig. 9, the ENM-BD computational cost has a marked linear scaling with protein size, owing to the absence of explicit solvent in the simulation and to the implementation of a neighbor-list approach to compute elastic interactions^34^. The few deviations from this trend correspond to particularly compact conformations, whose higher density of contacts results in a larger number of interactions. Together, these results indicate that ENM-BD is scalable to proteome-wide applications, to enable large-scale investigations of dynamics and flexibilities across hundreds of thousands of proteins at a computational cost that is several orders of magnitude lower than atomistic MD.

**Figure 9.**
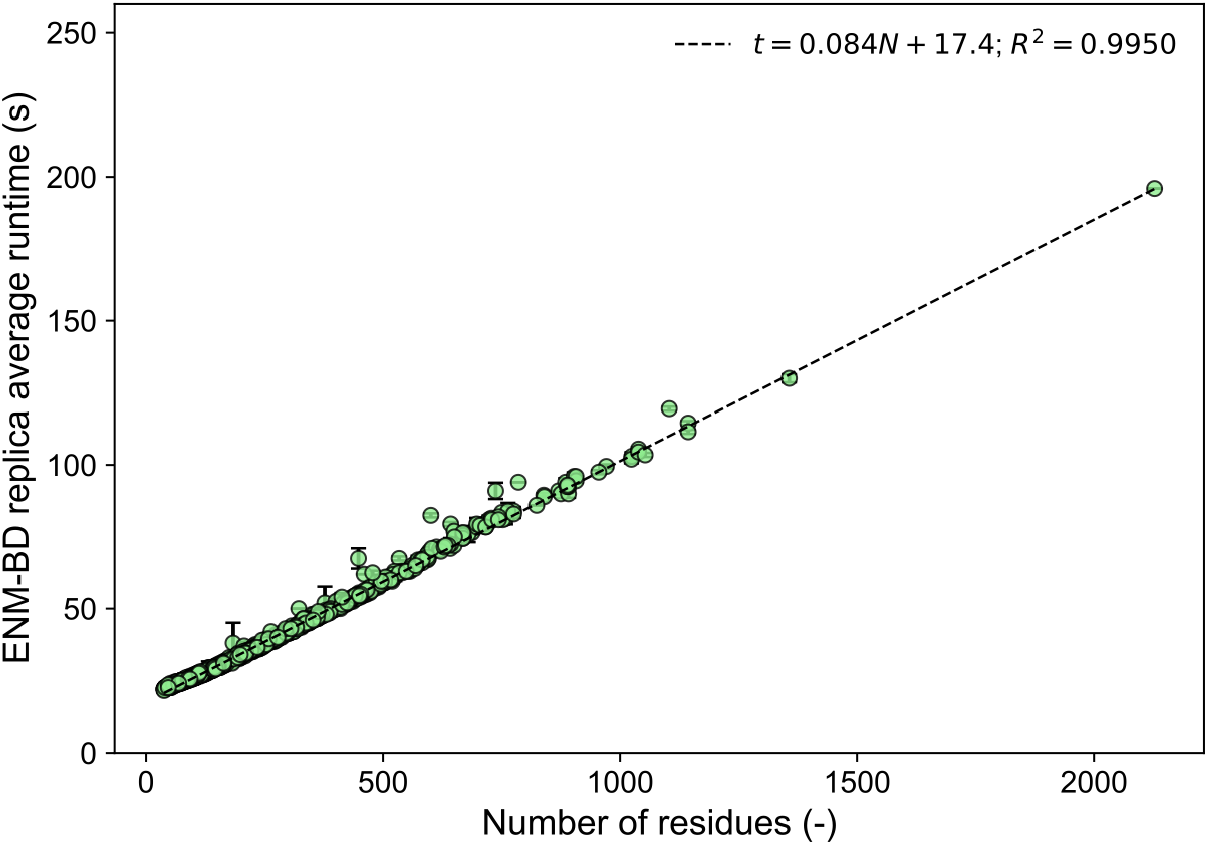
ENM-BD simulations are extremely fast and scale linearly with protein size. Each point represents the average wall-clock time required to simulate a 1 ns ENM-BD trajectory for a single protein (mean ± standard deviation over three independent replicas) from the ATLAS dataset, using a Linux workstation with 16 CPU threads. Running times are plotted as a function of protein size (number of residues). The equation of the fitting line, together with the *R*^*2*^ correlation value, is reported in the legend.

Despite the agreement with MD and NMR ensembles and its computational efficiency, it is important to note that ENM-BD retains intrinsic limitations inherited from the underlying network model. First, the current version of the model is completely chemistry-agnostic, as neither amino acid-specific interactions nor explicit solvent effects are considered. As a result, conformational changes driven by specific chemical interactions, charge interactions, ligand binding, or solvent-mediated effects cannot be modelled. Second, the ENM relies on harmonic potentials centered on the initial structure, making the starting conformation an energy minimum by construction^16^. Consequently, although stochastic Brownian forces allow the protein to explore alternative conformations, the trajectories tend to remain biased toward the vicinity of the initial state. Large-scale conformational transitions can thus only emerge if many native contacts are disrupted during the simulation. While this limits the exploration of distant regions of the conformational landscape, it also partially explains the good agreement observed with fluctuation patterns and essential motions from 100 ns-long MD and NMR, which typically do not account for dramatic, large-scale conformational transitions. Including additional terms to the underlying ENM, such as electrostatic and hydrophobic/polar interactions, might improve the description of chemically driven conformational changes, potentially allowing to enhance the sampling of the conformational space. Yet, this will require extensive efforts in parametrizing such new interactions without jeopardizing the robustness and computational efficiency of the current framework.

Finally, we must highlight that BD simulations are completely unaware of normal modes. Unlike NMA, which deterministically computes linear motions by diagonalizing the Hessian matrix, BD generates non-linear trajectories by integrating the Langevin equations with stochastic contributions driven by random initial velocities and arising from thermal noise (see Equation 6). In addition, BD explicitly incorporates environmental parameters such as temperature *T* and friction timescale *τ*, whereas NMA depends solely on the topology of the elastic network. Yet, the fluctuations generated by ENM-BD exhibit an almost perfect correlation (∼95%) with those predicted by ENM-NMA, and the dominant modes emerging from BD trajectories align remarkably well with the corresponding normal modes (> 80%; Fig. 7). Similar observations were reported by Gō and co-workers in the context of atomistic MD and atomistic NMA^77^. In that case, the agreement was lower due to the large anharmonic effects in the atomistic dynamics^77^, whereas in the present framework the use of a purely elastic (harmonic) potential leads to a much closer correspondence between the BD- and NMA-derived motions. These results suggest that the conformational dynamics emerging from BD trajectories, consistent with those observed in MD simulations and NMR ensembles, is already encoded in the three-dimensional protein topology, which is naturally captured by ENM and described by normal modes. Yet, BD provides a useful and efficient way to move beyond the linear vector approximations of NMA and generate richer structural ensembles (Fig. 8).

## Author contribution

Conceptualization: D.S., B.H.L., L.O.; Methodology and model development: D.S., E.R., M.F.; Software implementation: D.S., E.R., M.F.; Data curation and computational analysis: D.S., E.R., M.F.; Validation and benchmarking: D.S., E.R., M.F., B.H.L.; Writing – original draft: D.S.; Writing – review and editing: D.S., E.R., M.F., B.H.L., L.O.; Supervision: D.S., L.O.; Funding acquisition: L.O.

## Funding and acknowledgments

L.O. acknowledges financial support from Cancerfonden Junior Investigator Award (CF 21 0305 JIA) and Project Grants (CF 21 1471 Pj, CF 24 3801 Pj) as well as Vetenskapsrådet Starting Grant (VR 2021-02248) and Karolinska Institutet. D.S. acknowledges financial support from Cancerfonden Postdoctoral Fellowship (CF 24 0908 PT).

## Data availability

The edENM^mod^-BD and eBDIMS2^mod^ codes to generate unbiased and biased trajectories are available at https://github.com/domenicoscaramozzino/ENM_BD. All the main data presented in the paper, including complete BD ensembles and performance metrics, are available at https://doi.org/10.6084/m9.figshare.33965158.

## Notes

https://github.com/domenicoscaramozzino/ENM_BD

https://doi.org/10.6084/m9.figshare.33965158

### Competing Interest Statement

The authors have declared no competing interest.

