## Supporting Information for "Leveraging elastic networks in a coarse-grained brownian simulation framework for protein conformational dynamics"

**Supplementary Table 1.** Mean and standard deviation of consecutive $C_\alpha$-$C_\alpha$ distances measured across the 1,938 monomeric proteins of the ATLAS dataset[1]. Values in parentheses indicate the Euclidean distance between each ENM-BD distribution and the MD distribution.

| Model | $C_\alpha$-$C_\alpha$ distance (Å) |
|---|---|
| ANM-BD | 3.87 ± 0.36 (0.231) |
| edENM-BD | 3.83 ± 0.10 (0.092) |
| $edENM^{mod}$-BD | 3.83 ± 0.07 (0.005) |
| MD | 3.83 ± 0.07 |

**Supplementary Table 2.** Mean and standard deviation of MolProbity scores[2] across the 1,938 monomeric proteins in the ATLAS dataset. Stereochemical quality was evaluated for the final frames of MD and ENM-BD simulations after reconstruction with cg2all[3]. "MD + cg2all" denotes MD conformers stripped to $C_\alpha$ atoms and subsequently reconstructed with cg2all. "$edENM^{mod}$-BD + cg2all + minimization" denotes reconstructed $edENM^{mod}$ ENM-BD conformations following a short energy minimization to remove reconstruction-induced artifacts.

| Model | MolProbity score (-) |
|---|---|
| MD | 1.388 ± 0.279 |
| ANM-BD + cg2all | 3.350 ± 0.216 |
| edENM-BD + cg2all | 2.681 ± 0.289 |
| $edENM^{mod}$-BD + cg2all | 2.330 ± 0.305 |
| MD + cg2all | 2.229 ± 0.287 |
| $edENM^{mod}$-BD + cg2all + minimization | 1.278 ± 0.315 |

**Supplementary Table 3.** Mean and standard deviation of agreement metrics between ENM-BD and atomistic MD across the ATLAS dataset: correlations of residue fluctuations (RMSF PCC), maximum overlaps of principal components ($O_{max}$), and PC subspace similarity ($RMSIP_2$).

| Model | RMSF PCC (-) | $O_{max}$ (-) | $RMSIP_2$ (-) |
|---|---|---|---|
| ANM-BD | 0.762 ± 0.163 | 0.698 ± 0.120 | 0.559 ± 0.202 |
| edENM-BD | 0.799 ± 0.163 | 0.726 ± 0.120 | 0.592 ± 0.201 |
| $edENM^{mod}$-BD | 0.795 ± 0.156 | 0.708 ± 0.118 | 0.581 ± 0.197 |

**Supplementary Table 4.** Mean and standard deviation of consecutive $C_\alpha$-$C_\alpha$ distances measured across the 6,704 monomeric proteins in the NMR dataset. The value in parentheses indicates the Euclidean distance between the ENM-BD distribution and the corresponding NMR distribution.

| **Model** | **$C_\alpha$-$C_\alpha$ distance (Å)** |
|---|---|
| edENM$^{mod}$-BD | 3.81 ± 0.05 (0.073) |
| NMR | 3.81 ± 0.07 |

**Supplementary Table 5.** Mean and standard deviation of MolProbity scores across the 6,704 monomeric proteins in the NMR dataset. "NMR + cg2all" denotes NMR models stripped to $C_\alpha$ atoms and subsequently reconstructed with cg2all.

| **Model** | **MolProbity score (-)** |
|---|---|
| NMR | 3.035 ± 0.876 |
| edENM$^{mod}$-BD + cg2all | 2.824 ± 0.384 |
| NMR + cg2all | 2.697 ± 0.426 |
| edENM$^{mod}$-BD + cg2all + minimization | 1.704 ± 0.449 |

**Supplementary Table 6.** Mean and standard deviation of agreement metrics between ENM-BD and NMR ensembles, including correlations of residue fluctuations (RMSF PCC), maximum overlaps of principal components ($O_{max}$), and PC subspace similarity ($RMSIP_2$).

| **Model** | **RMSF PCC (-)** | **$O_{max}$ (-)** | **$RMSIP_2$ (-)** |
|---|---|---|---|
| edENM$^{mod}$-BD | 0.816 ± 0.176 | 0.689 ± 0.137 | 0.602 ± 0.195 |

**Supplementary Table 7.** Mean and standard deviation of agreement metrics between ENM-NMA and ENM-BD for the MD and NMR dataset.

| **Dataset** | **RMSF PCC (-)** | **$O_{max}$ (-)** | **$RMSIP_2$ (-)** |
|---|---|---|---|
| ATLAS | 0.961 ± 0.069 | 0.939 ± 0.061 | 0.889 ± 0.123 |
| NMR | 0.936 ± 0.094 | 0.900 ± 0.084 | 0.828 ± 0.144 |

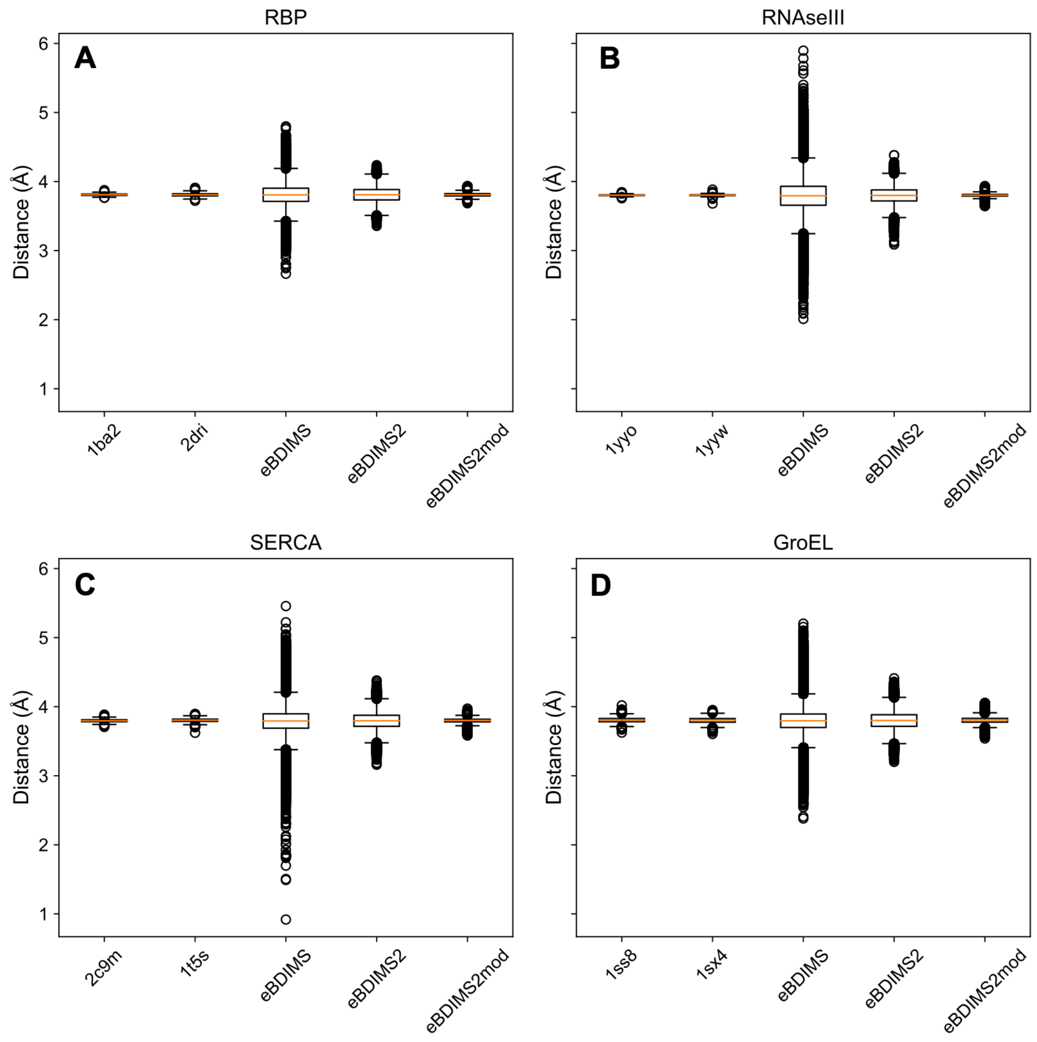


**Supplementary Figure 1. edENM$^{mod}$ preserves the local backbone geometry during large-scale conformational transitions.** Distribution of consecutive $C_\alpha$-$C_\alpha$ distances for four benchmark proteins previously analyzed in Scaramozzino et al.[4]: (A) ribose-binding protein (RBP; PDB IDs: 1ba2 and 2dri); (B) RNA endonuclease III (RNaseIII; 1yyo and 1yyw); (C) sarcoplasmic/endoplasmic reticulum Ca2+ ATPase1 (SERCA; 2c9m and 1t5s); (D) GroEL chaperonin heptamer (1ss8 and 1sx4). Boxplots compare the consecutive $C_\alpha$-$C_\alpha$ distances in the experimental end-state structures with those sampled along transition pathways generated by eBDIMS[5], eBDIMS2[4], and eBDIMS2 employing edENM$^{mod}$. Boxes represent the interquartile range (25$^{th}$ and 75$^{th}$ percentiles), the central line indicates the median, whiskers extend to minimum and maximum non-outlier values, and outliers are shown as dots.

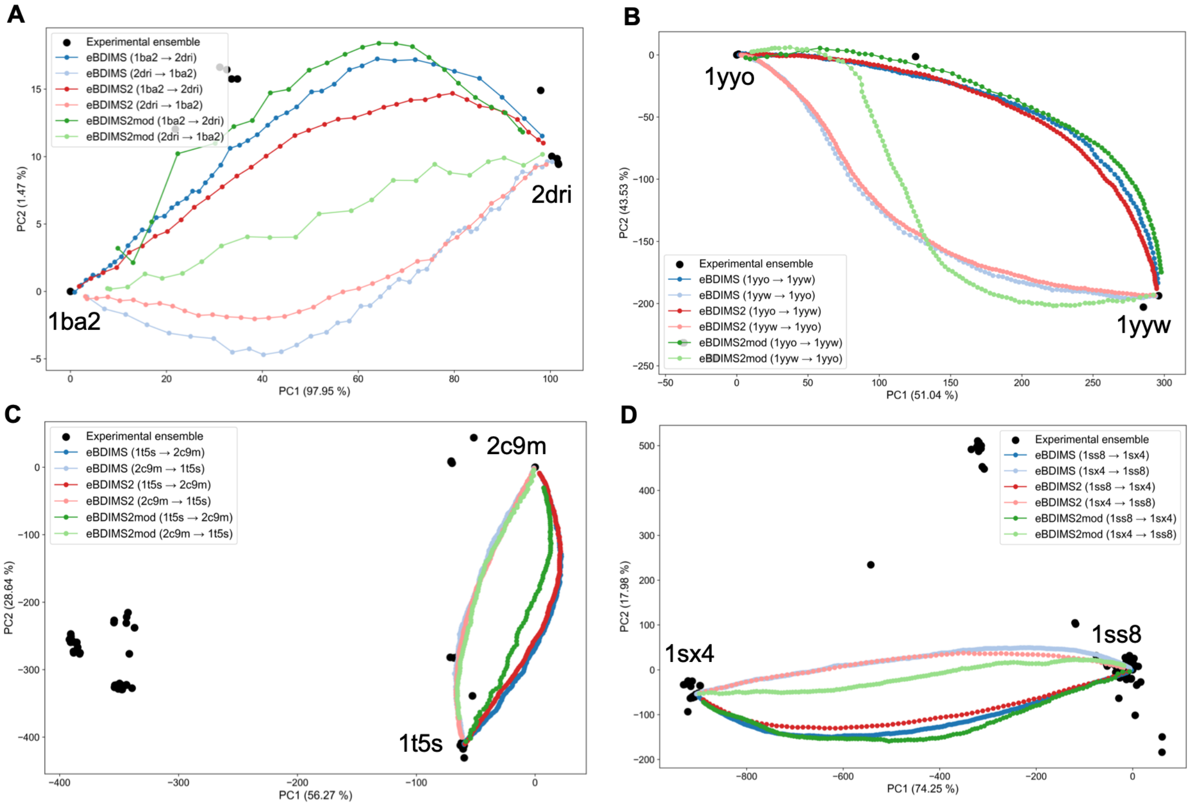


**Supplementary Figure 2. The modified elastic network does not drastically modify transition pathways predicted by eBDIMS.** Transition pathways between open and closed conformations generated by eBDIMS[5] (blue), eBDIMS2[4] (red), and eBDIMS2 employing edENM$^{mod}$ (green) for (A) RBP, (B) RNaseIII, (C) SERCA, and (D) GroEL chaperonin heptamer. The trajectories have been projected in the 2D space of the lowest PCs of the experimental ensembles as in Scaramozzino et al.[4]. Other than the edENM$^{mod}$ variant, eBDIMS2$^{mod}$ implementation also employs updated BD parameters ($\Delta t = 2$ fs; $\tau = 2.5$ ps) compared to the previous eBDIMS and eBDIMS2 versions ($\Delta t = 1$ fs; $\tau = 0.004$ ps).

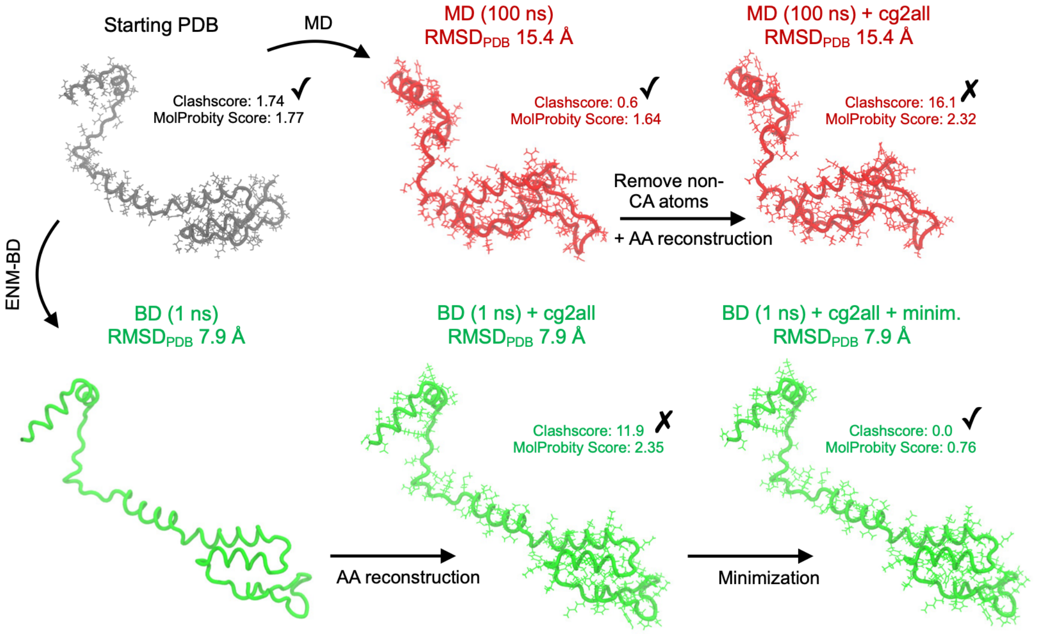


**Supplementary Figure 3. Representative example illustrating the effect of CG-to-all-atom reconstruction on stereochemistry quality.** The starting X-ray structure of the secreted effector protein SptP (PDB: 1jyo, chain E) has excellent stereochemistry (MolProbity score = 1.77; clash score = 1.74). Top row (red): the final frame of a 100 ns atomistic MD trajectory (replica 3; $C_\alpha$ RMSD ~ 15 Å from the starting structure) retains high stereochemical quality (MolProbity = 1.64; clash score = 0.6). However, stripping the structure to $C_\alpha$ atoms and subsequently reconstructing with cg2all decreases the MolProbity score to 2.32 (clash score = 16.1), demonstrating that the reconstruction procedure itself can induce stereochemical defects. Bottom row (green): the final frame of a 1 ns ENM-BD simulation with edENM$^{mod}$ ($C_\alpha$ RMSD ~ 8 Å) after cg2all reconstruction exhibits a comparable MolProbity score (2.35; clash score = 11.9). A brief atomistic energy minimization restores the stereochemistry (MolProbity = 0.76; clash score = 0.0), while preserving the sampled coarse-grained conformation (RMSD ~ 0.2 Å with respect to non-minimized conformation). Structures were rendered with VMD[6].

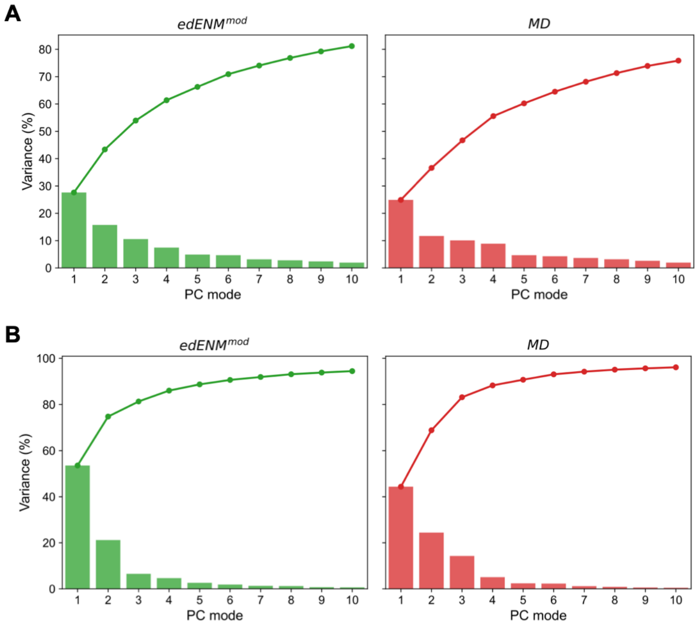


**Supplementary Figure 4. Comparison of PC variance distributions for ENM-BD and MD ensembles.** (A) *D. Melanogaster* caprin homolog (PDB: 6bk4, chain: A); (B) DUF5617 domain-containing protein from *L. pneumophila* (4wrp, A). Variances are shown for ENM-BD (left, green) and MD (right, red) ensembles. Histogram bars represent the variance associated with each PC vector, while solid lines indicate the cumulative variance.

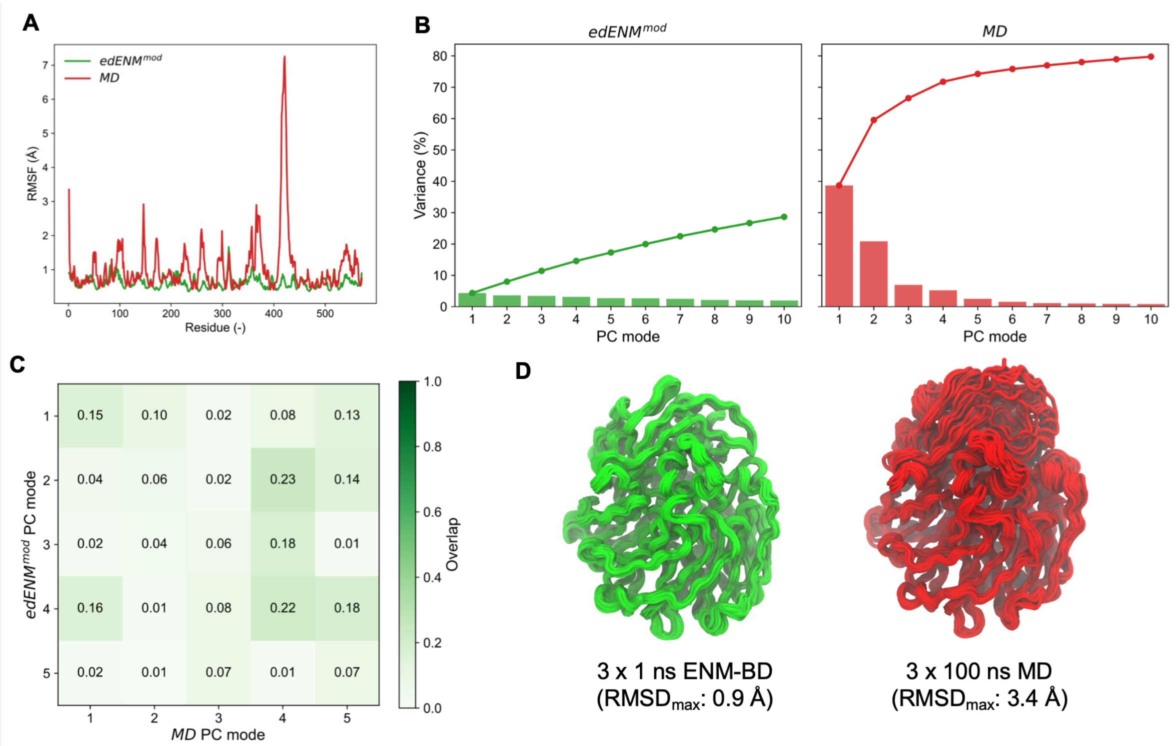


**Supplementary Figure 5. A representative case of poor agreement between ENM-BD and atomistic MD.** *M. Methylotrophus* methanol dehydrogenase subunit 1 (PDB: 2ad6, chain: A). (A) Comparison of RMSFs obtained from ENM-BD simulations (green) and MD (red). (B) Distribution of variances along the dominant PC modes from ENM-BD (left) and MD (right). (C) Overlap matrix between the five leading PC vectors from ENM-BD and MD, with values ranging from 0 (white) to 1 (dark green). (D) $C_\alpha$-tube representations of the conformational ensembles sampled by ENM-BD (green) and MD (red).

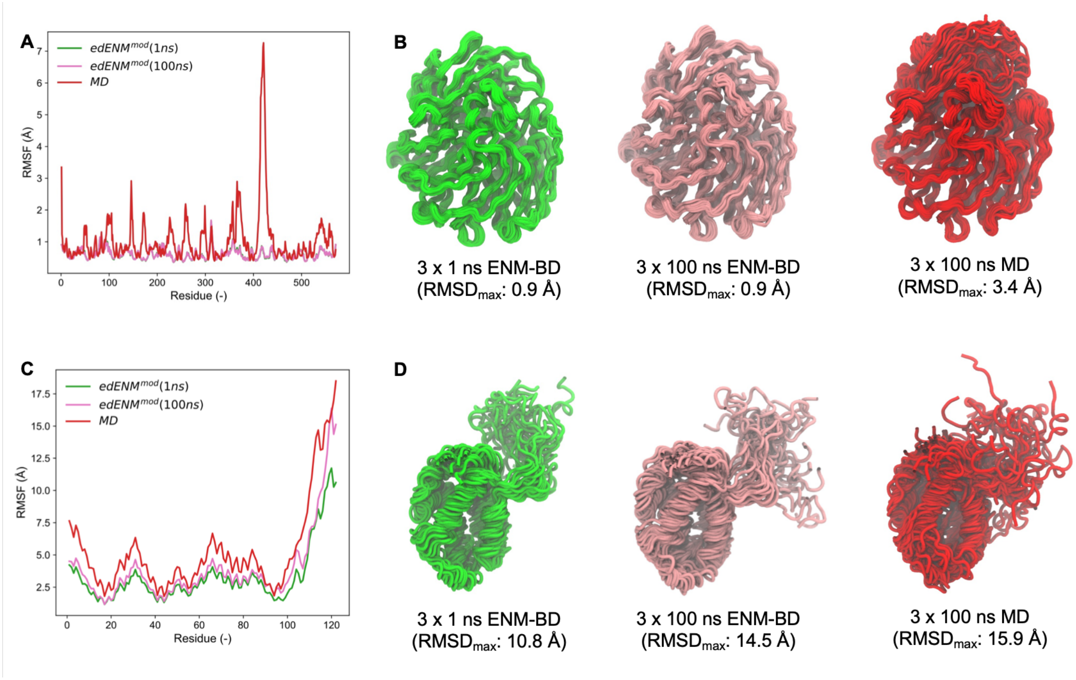


**Supplementary Figure 6. Effect of simulation length on ENM-BD conformational sampling.** (A-B) *M. Methylotrophus* methanol dehydrogenase subunit 1 (2ad6, A). (C-D) *L. pneumophila* DUF5617 domain-containing protein (PDB: 4wrp, chain: A). (A,C) Comparison of RMSF profiles obtained from atomistic MD (red), short ENM-BD (1 ns; green), and longer ENM-BD simulations (100 ns; pink). (B, D) $C_\alpha$-tube representations of conformational ensembles generated by MD (red), short ENM-BD (green) and longer ENM-BD (pink), illustrating the effect of increasing simulation time on the sampled conformations.

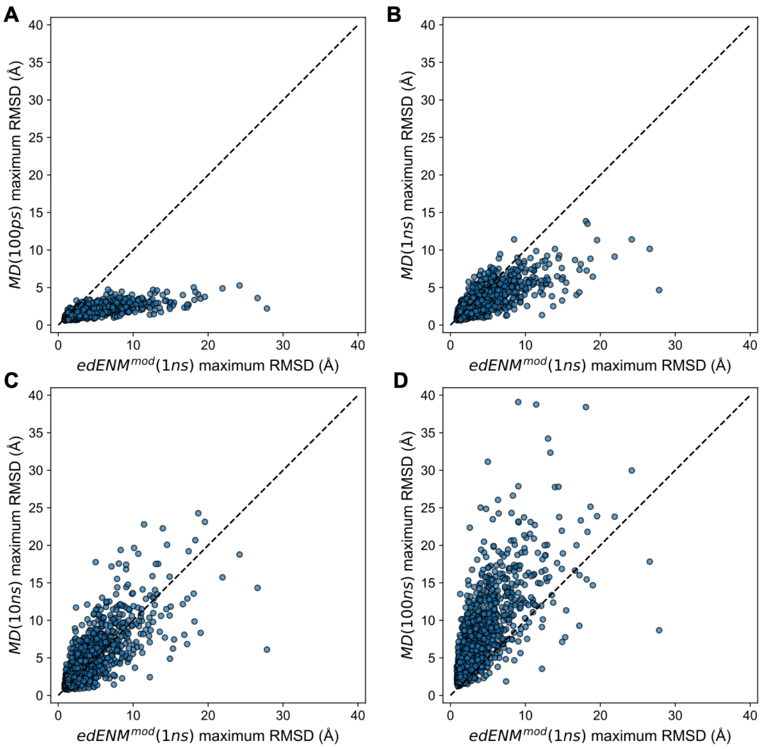


**Supplementary Figure 5. Comparison between 1 ns-long edENM$^{mod}$-BD and MD sampling cut at different timescales.** Each point represents a protein in the ATLAS dataset and reports the maximum RMSD from the initial X-ray structure obtained from 1 ns-long BD simulations (X-axis) and atomistic MD simulations (Y-axis) at different timescales: (A) 100 ps; (B) 1 ns; (C) 10 ns; (D) 100 ns.

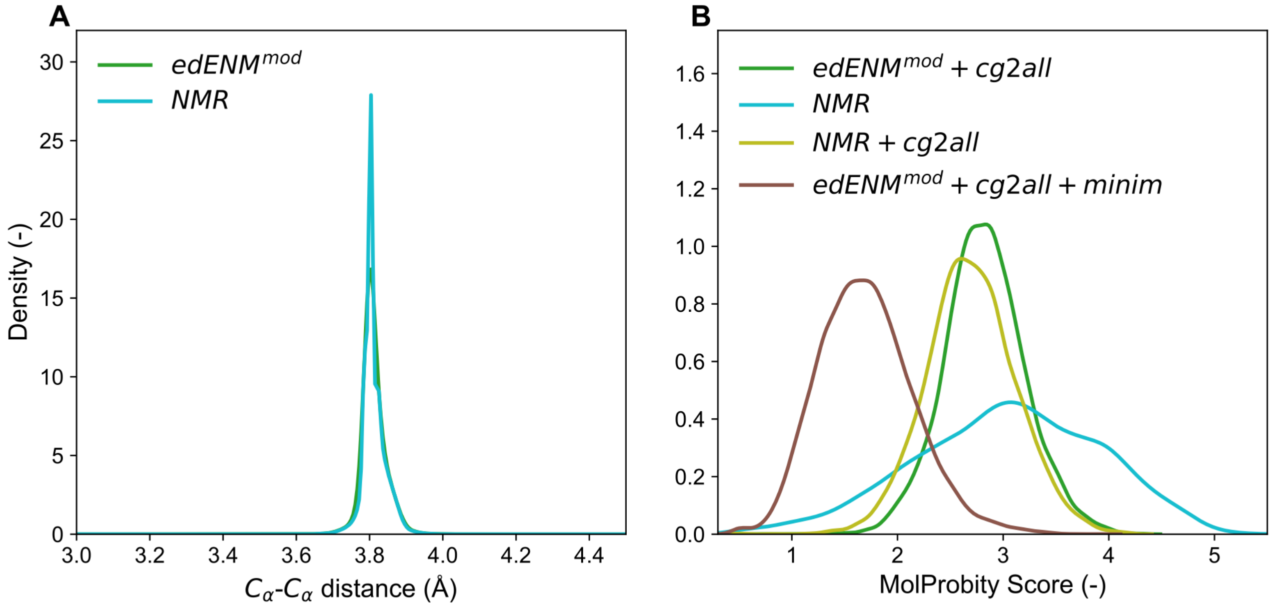


**Supplementary Figure 6. Stereochemical quality of ENM-BD conformers across the NMR dataset.** (A) Distribution of consecutive $C_\alpha$-$C_\alpha$ distances in NMR models (cyan) and ENM-BD simulations performed with edENM$^{mod}$ (green). (B) MolProbity scores of NMR models (cyan), edENM$^{mod}$ ENM-BD conformers after cg2all reconstruction (green), NMR conformers stripped to $C_\alpha$ atoms and reconstructed with cg2all (lime yellow), and reconstructed edENM$^{mod}$ conformers following a short energy minimization (brown).

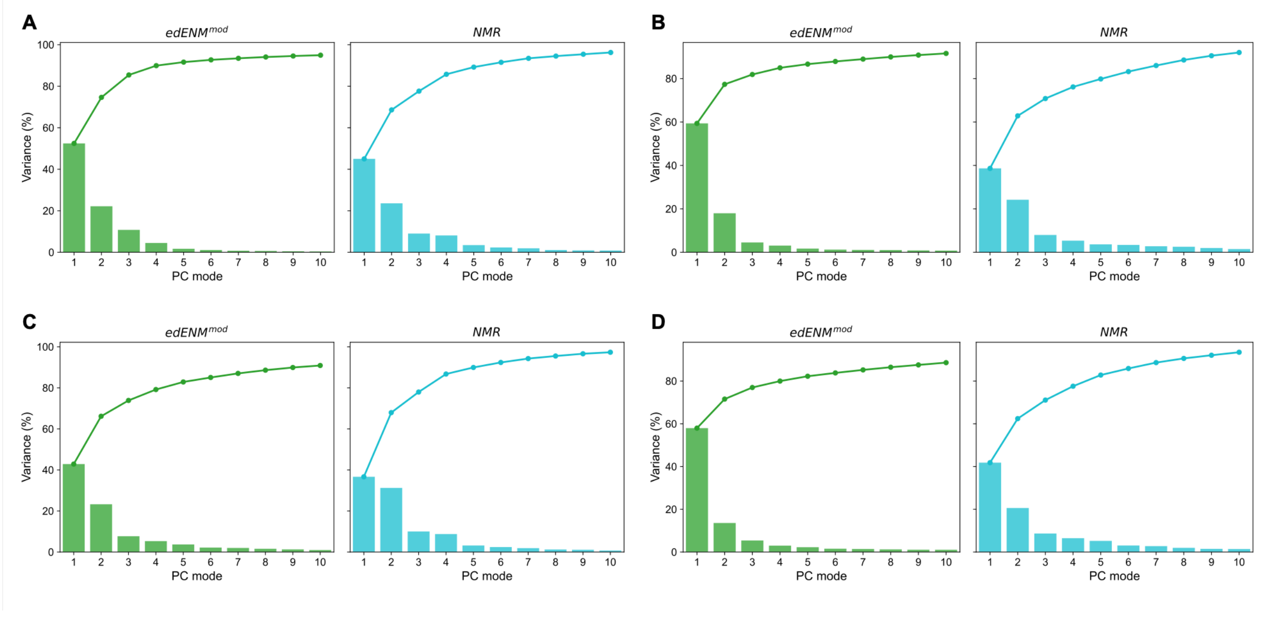


**Supplementary Figure 7. Comparison of PC variance distributions for ENM-BD and NMR ensembles with remarkable ensemble agreement.** (A) C-terminal domain of *H. Sapiens* Lamin-B2 (PDB: 2lll, chain: A); (B) synthetic *de novo* design of ferredoxin-like fold protein (2kl8, A); (C) fibronectin type III domain of *H. Sapiens* tenascin X (2cui, A); (D) DUF971 domain-containing protein from *P. Aeruginosa* (2l6p, A). Variances are shown for ENM-BD (left, green) and NMR (right, cyan) ensembles. Histogram bars represent the variance associated with each PC vector, while solid lines indicate cumulative variances.

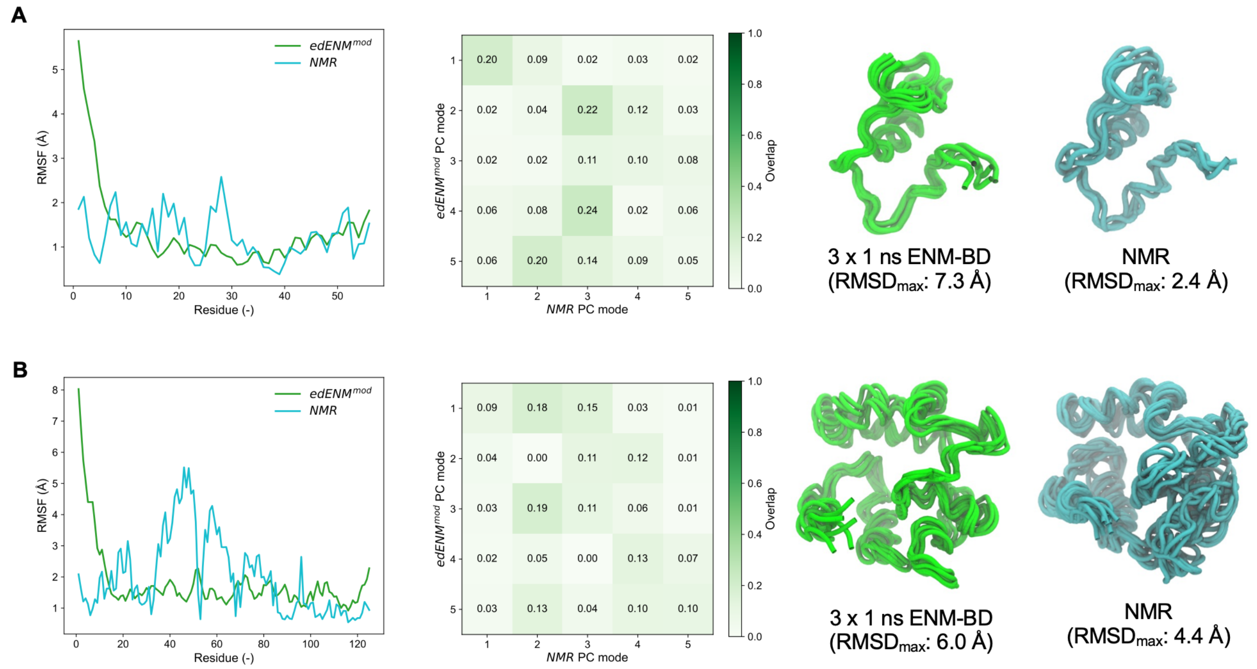


**Supplementary Figure 8. Two representative cases of limited agreement between ENM-BD and NMR.** (A) *B. Taurus* proteinase inhibitor IIA (PDB: 1bus, chain: A); (B) *H. Sapiens* vesicle-associated membrane protein-associated protein B/C (2mdk, A). Left panels compare $C_\alpha$ RMSF profiles obtained from ENM-BD (green) and NMR ensembles (cyan). Middle panels show overlap matrices between the five leading principal components of the ENM-BD and NMR ensembles. Right panels display $C_\alpha$-tube representations of the conformational ensembles by ENM-BD (green) and NMR (cyan).

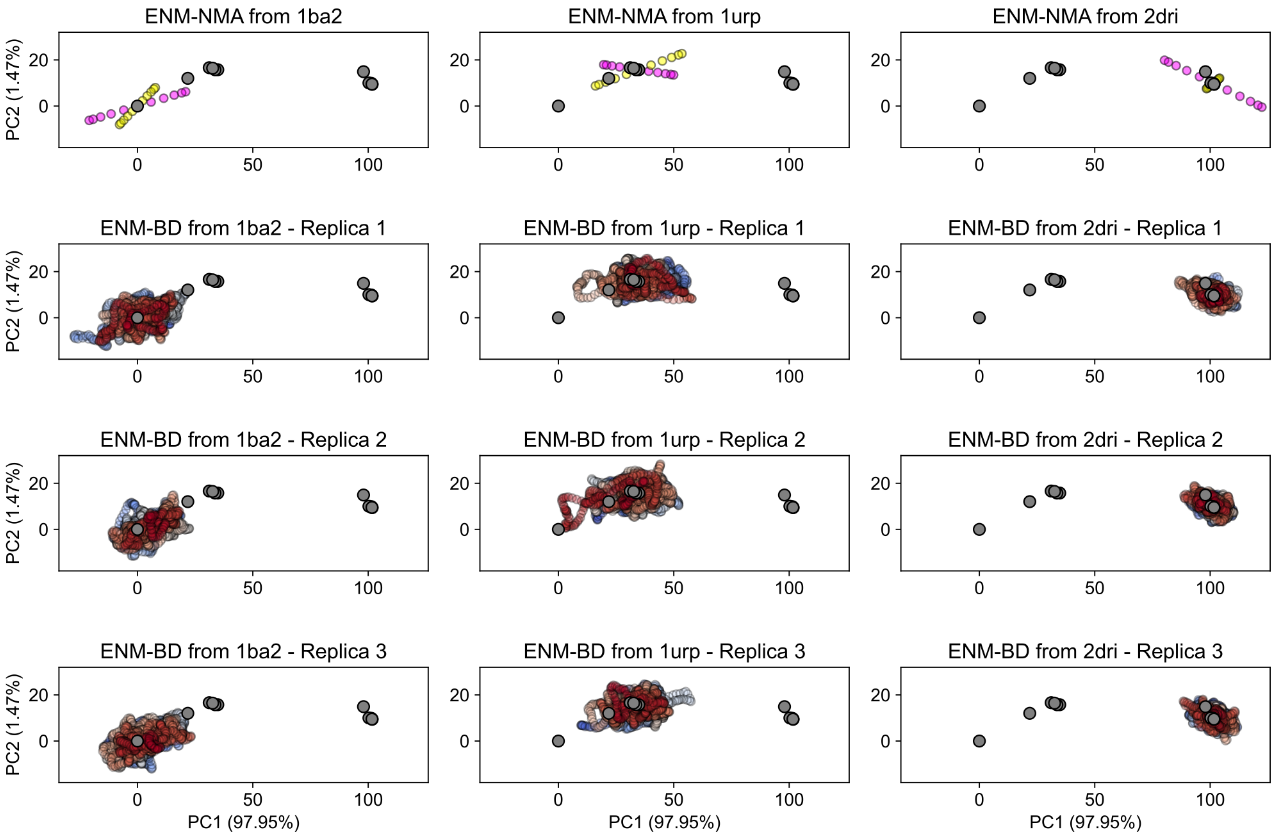


**Supplementary Figure 9. Projection of conformations generated by ENM-NMA and ENM-BD onto the experimental PCA space of ribose-binding protein (RBP).** The first row shows the projection of conformations generated by linear extrapolation along the first (NM1, magenta) and second (NM2, yellow) normal modes, respectively. The last three rows display projections of conformations sampled by three independent 1 ns BD replicas. Point colors indicate simulation progress, ranging from blue (initial frame) to red (final frame). Columns correspond to calculations initiated from the open (PDB: 1ba2), intermediate (1urp), and closed (2dri) conformations. Grey points represent the experimental X-ray structures of RBP[4,7].
